# *Papaver S*-determinants trigger an integrated network of mitochondrially derived ROS and disruption of energy metabolism in incompatible pollen tubes

**DOI:** 10.1101/2025.06.26.661469

**Authors:** Ludi Wang, An-Shan Hsiao, José Carli, Ali Raza, Zongcheng Lin, Dominique Arnaud, Julia M. Davies, Vernonica E. Franklin-Tong, Nicholas Smirnoff, Maurice Bosch

## Abstract

Many plants use self-incompatibility (SI) mechanisms to prevent inbreeding by inhibiting pollen tube growth in “self” pistils. In *Papaver rhoeas*, SI is triggered by allele-specific interaction between the pollen and pistil *S*-determinants, activating a Ca^2+^-dependent signalling network that leads to rapid reactive oxygen species (ROS) production and eventual programmed cell death (PCD) in incompatible pollen tubes. Expression of the *Papaver* pollen *S*-determinant (PrpS) in *Arabidopsis thaliana* recapitulates *Papaver* SI when challenged with the cognate pistil ligand (PrsS). Using roGFP2-Orp1, a genetically encoded hydrogen peroxide (H_2_O_2_) sensor, combined with measurements of mitochondrial metabolism we have revealed a complex SI response. Within minutes, elevated cytosolic Ca^2+^ ([Ca^2+^]_cyt_) and cytosolic acidification converge to trigger mitochondrial H_2_O_2_ production (with consequent cytosolic and plastid H_2_O_2_ increase), membrane depolarisation, decreased respiration rate, and ATP depletion. In parallel, oxidative inactivation of cytosolic GAPDH inhibits glycolytic flux, resulting in decreased TCA cycle intermediates and providing a feedback loop to enhance mitochondrial disruption. Preceding production of mitochondrially-sourced ROS, we show that SI rapidly arrests pollen tube growth *via* inactivation of plasma membrane localised NADPH oxidase (RBOH) mediated superoxide production. Our findings provide new insights into how different ROS signatures from NADPH oxidase and mitochondria drive distinct processes. We demonstrate that mitochondrial disruption is an early and central event in the SI-response, likely driven by positive feedback loops involving altered Ca^2+^ signalling, cytosolic pH and ROS. Our findings provide novel insights into how SI rapidly disrupts energy metabolism in pollen tubes through interconnected signalling networks centred on mitochondria prior to PCD.

## Introduction

Self-incompatibility (SI) is a pollen-pistil recognition system utilized by many flowering plants to prevent self-fertilization and maintain genetic diversity. This system is governed by tightly linked, polymorphic *S*-determinants expressed in the pollen and the pistil, which regulate mating compatibility and prevent inbreeding. In *Papaver rhoeas* (poppy), the *S*-determinants are the stigma-expressed S-protein (PrsS), which is secreted and acts as the ligand, and its pollen-specific transmembrane receptor (PrpS) (Foote et al., 1994; Wheeler et al., 2009). Interaction between these two proteins triggers a rapid growth arrest of incompatible pollen tubes mediated by a cytosolic free calcium ([Ca^2+^]_cyt_)-dependent signalling network that includes increases in reactive oxygen species (ROS), cytosolic acidification, and rapid ATP depletion (Wilkins et al., 2011; Wilkins et al., 2015; Wang et al., 2022; Bosch and Franklin-Tong, 2024). These events trigger programmed cell death (PCD) of incompatible pollen tubes several hours later (Thomas and Franklin-Tong, 2004; Wang et al., 2019). Understanding how these molecular and cellular events coordinate pollen compatibility and control cell survival or death provides key insights into plant reproductive regulation and offers broader implications for manipulating crop fertility and breeding strategies.

SI in *Papaver* stimulates dramatic increases in ROS levels throughout the entire pollen tube, as demonstrated by the non-specific ROS-sensitive probe chloromethyl-2’7’-dichlorodihydrofluorescein diacetate acetyl ester (CM-H_2_DCFDA) (Hempel et al., 1999; Wilkins et al., 2011). This distinctive pattern of ROS accumulation raises important questions - what are the sources and targets of ROS during SI? To investigate these questions, a means of measuring ROS with greater spatial and temporal resolution is required. Recent advances in fluorescent biosensors, such as the H_2_O_2_-specific probe roGFP2-Orp1, now allow for precise and dynamic measurements of H_2_O_2_ levels (Nietzel et al., 2019). This sensor has been effectively used to study H_2_O_2_ production during pattern-triggered immunity (PTI) responses (Arnaud et al., 2023) and mitochondrial retrograde signalling (Khan et al., 2024) in *Arabidopsis*. Notably, the *Papaver* SI system has been functionally transferred to *Arabidopsis thaliana* (de Graaf et al., 2012; Lin et al., 2015, Wang et al., 2020; Wang et al., 2022). This model system, utilizing pollen tubes expressing PrpS, combined with the genetically encoded fluorescent probe roGFP2-Orp1, offers a powerful platform to investigate SI-induced subcellular H_2_O_2_ dynamics in greater depth using live-cell imaging approaches.

Pollen tube growth is an energy-intensive process, requiring high ATP levels to fuel the cellular machinery essential for tip growth (Rounds et al., 2011). We recently established that SI triggers a rapid and dramatic reduction in ATP levels (Wang et al., 2022) and that SI-induced ROS triggers irreversible oxidation of multiple enzymes, including those involved in energy metabolism (Haque et al., 2020). Reduced ATP synthesis implicates respiratory metabolism and mitochondria as early targets in the SI-induced PCD pathway. Supporting this, our previous studies revealed that SI triggers a rapid release of cytochrome *c* (Thomas and Franklin-Tong, 2004) and leads to significant alterations in mitochondrial morphology (Geitmann et al., 2004) in *Papaver* pollen tubes.

Cytosolic pH ([pH]_cyt_) and [Ca^2+^]_cyt_ also play crucial roles in pollen tube growth. Under normal conditions, pollen tubes exhibit an apical gradient of both [pH]_cyt_ and [Ca^2+^]_cyt_, which mirror the oscillatory growth pattern characteristic of tip-growing cells (Holdaway-Clarke & Hepler, 2003; Hepler et al., 2006). However, during SI, these tip-based ion gradients collapse, accompanied by elevated [Ca^2+^]_cyt_ (Franklin-Tong et al., 1993) and dramatic cytosolic acidification (Bosch and Franklin-Tong, 2007; Wilkins et al., 2015; Wang et al., 2022), which cause cessation of growth and activation of PCD.

ROS, primarily superoxide generated *via* electron transport and oxidase enzymes, and its dismutation product hydrogen peroxide (H_2_O_2_) play a crucial role in plant development and stress responses (Smirnoff and Arnaud, 2019), yet excessive ROS accumulation leads to PCD (Mittler et al., 2022). In plant cells that utilize polar growth, such as pollen tubes and root hairs, tip-localized superoxide production mediated by plasma membrane NADPH oxidases (RBOHs), is critical for sustained tip growth (Foreman et al., 2003; Potocký et al., 2007; Kaya et al., 2014; Smirnoff and Arnaud, 2019). Recent evidence suggests that the intracellular origin of ROS plays a pivotal role in determining their signalling functions (Noctor & Foyer, 2016; Mittler et al., 2022; Arnaud et al., 2023; Khan et al., 2024), adding further complexity to their regulation and underscoring the need for precise tools to dissect their roles.

In this study, we show that SI rapidly inhibits tip-localised superoxide production, concomitant with the inhibition of tip growth. Using the H_2_O_2_-specific sensor roGFP2-Orp1, we demonstrate increases in probe oxidation in the cytosol, mitochondria and plastids. This reveals that ROS signals regulating growth and SI arise from distinct subcellular origins and exhibit unique spatial-temporal signatures. We also show that increases in [Ca^2+^]_cyt_ and cytosolic acidification contribute to SI-induced mitochondrial dysfunction and H_2_O_2_ generation. Moreover, SI triggers a collapse in mitochondrial membrane potential, leading to reduced respiration and decreased levels in TCA cycle and glycolysis intermediates. Together, our findings provide critical insights into the interplay between [Ca^2+^]_cyt_, cytosolic pH, H_2_O_2_ and mitochondria during SI, illustrating how these signals interact and converge to disrupt energy metabolism in pollen tubes.

## Results

### SI induces rapid inactivation of pollen tube tip-localised ROS production

In our experimental system, SI was induced by applying recombinant PrsS_1_ protein to *in vitro* growing Arabidopsis pollen tubes expressing the *Papaver PrpS_1_* gene (de Graaf et al., 2012). Because SI causes increased ROS production in the pollen tube during SI (Wilkins et al., 2011), we measured production of tip-localised superoxide. Tip growth is dependent on ROS generated by NADPH oxidase activity (Potocký et al., 2007; Boisson-Dernier 2013; Lassig et al., 2014; Kaya et al., 2015). Apoplastic superoxide production, driven by NADPH oxidases at the tip of growing pollen tubes, can be visualised by the reduction of nitroblue tetrazolium (NBT) to an insoluble blue formazan. NBT reduction is largely apoplastic and is abolished by knockdown of NADPH oxidase expression (Potocký et al., 2007). In untreated Arabidopsis pollen tubes, tip-localised formazan staining was observed (**Figure 1A**), indicating superoxide production at the tip. Following SI-induction, tip-localised superoxide production was rapidly disrupted (**Figure 1A-B**, supplemental **Figure S1**). Within 5 minutes, the intensity of formazan staining was significantly reduced, and this reduction plateaued by 10-15 min (**Figure 1A-B**). A similar response was observed in *Papaver* pollen tubes, where the basal level of apoplastic superoxide was higher, leading to a more pronounced decrease after SI induction (Supplementary **Figure S1**). Inactivation of NADPH oxidase corresponds to the rapid inhibition of tip growth during SI (**Videos 1 and 2**). To establish whether this disruption of NADPH oxidase activity was a consequence or cause of SI-induced growth inhibition, we treated pollen tubes with 500 μM GdCl_3_ or 10 mM caffeine, which are known inhibitors of tip growth (Pierson et al., 1996; Geitmann et al., 2000). Within two minutes of treatment, we observed significant decreases in NBT staining (**Figure 1C-D**), confirming that inhibition of tip growth rapidly inhibits tip-localised NADPH oxidase activity. As NADPH oxidase activity is inhibited by SI, abolishing tip-localized ROS production, it is evident that superoxide produced by NADPH oxidase cannot be the source of the SI-induced H_2_O_2_ in incompatible pollen tubes.

**Figure 1.**
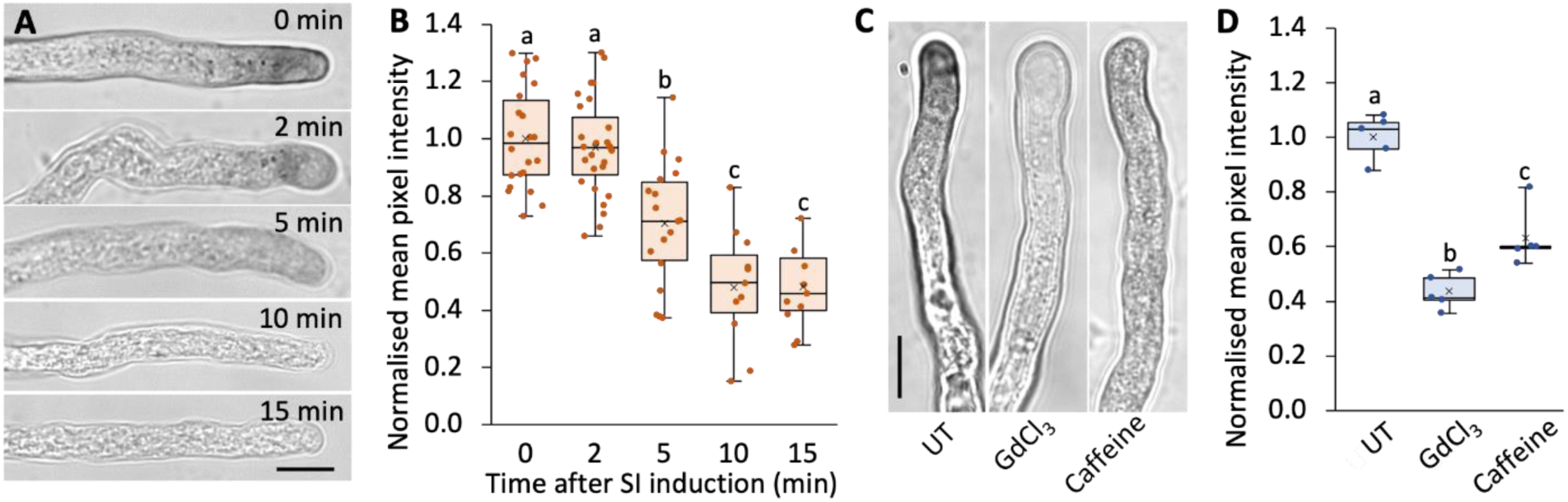
Inhibition of pollen tube tip-localised NADPH oxidase activity following SI induction. (A) Detection of superoxide production in Arabidopsis pollen tubes during the SI response using nitroblue tetrazolium (NBT) staining. Representative pollen tubes expressing PrpS_1_ were stained with 5 mg ml^-1^ NBT at 0, 2, 5, 10 and 15 minutes after SI induction. Scale bar = 10 μm. (B) Quantification of NBT staining intensity at pollen tube tips following SI induction. Reduced NBT staining corresponds to a decrease in pixel intensity. The mean pixel intensity at 0 minutes was normalised to 1 (n ≥ 10). NBT staining significantly decreased from 5 min after SI induction. (C) Reduced tip-localized superoxide production detected by NBT staining after 2 minutes of treatment with 500 μM GdCl_3_ or 10 mM caffeine. Scale bar = 10 μm. (D) Quantification of NBT staining intensity at pollen tube tips following treatment, with decreased pixel intensity indicating reduced superoxide production (n = 5). Different letters in (B) and (D) indicate significant (*P* < 0.05) differences based on Tukey’s test.

### Identifying the SI-induced intracellular ROS sources in pollen tubes

To investigate whether intracellular sources of ROS are involved in regulating SI, we examined ROS production in subcellular compartments during SI using transgenic Arabidopsis lines co-expressing PrpS_1_ and roGFP2-Orp1. An Orp1 cysteine residue is oxidised by H_2_O_2_ followed by a redox relay resulting in roGFP oxidation and a change in its fluorescence excitation spectrum, enabling ratiometric detection of its oxidation state (Nietzel et al., 2019). The roGFP2-Orp1 sensor, targeted to the cytosol, mitochondria, plastids, peroxisomes and nuclei, allowed compartment-specific H_2_O_2_ detection. Fluorescence imaging confirmed the subcellular localisation of the roGFP2-Orp1 sensor in growing pollen tubes (Supplementary **Figure S2A**). The oxidation of roGFP2-Orp1 by H_2_O_2_, measured ratiometrically as R_405/488_ (Nietzel et al., 2019), provides a quantitative readout of H_2_O_2_ dynamics. In untreated (UT) pollen tubes, roGFP2-Orp1 oxidation was relatively higher in mitochondria and peroxisomes, while the sensor was more reduced in the nuclei compared to the cytosol and plastids (Supplementary **Figure S2B**). The dynamic range (δ) of oxidation and reduction of roGFP2-Orp1 for each subcellular localisation was assessed by treating pollen tubes with 15 mM H_2_O_2_ or 10 mM dithiothreitol (DTT; Supplementary **Figure S2C-G**). The response of roGFP2-Orp1 to pollen tubes treated with H_2_O_2_ was more pronounced in the cytosol and mitochondria, while DTT treatment resulted in a greater reduction in plastids and peroxisomes compared to cytosol, mitochondria and nuclei. Despite nuclei (δ = 2.71) and peroxisomes (δ = 2.49) displaying a narrower dynamic range of roGFP2-Orp1 compared to the cytosol (δ = 3.65), mitochondria (δ = 5.09) and plastids (δ = 5.70), the probe characteristics are similar to previous reports and suitable for detecting changes in oxidation state (Nietzel et al., 2019; Ugalde et al., 2021; Arnaud et al., 2023).

To examine the changes in oxidation state across various subcellular compartments following SI induction, we performed ratio-imaging of roGFP2-Orp1 by sampling populations of pollen tubes (**Figure 2**). Probe oxidation is detected as an increase in the fluorescence excitation ratio (R_405/488_). We observed rapid increases in cytosolic and mitochondrial roGFP2-Orp1 oxidation, demonstrating an elevation in H_2_O_2_. In the cytosol, a significant increase in roGFP2-Orp1 oxidation was detectable as early as 10 minutes after SI induction compared to untreated pollen (**Figure 2A**). At 60 minutes, increases in roGFP2-Orp1 oxidation became more pronounced, increasing further to 2.3-fold by 2 hours after the SI induction (**Figure 2A**). In mitochondria, a significant increase in probe oxidation was also observed as early as 10 minutes after SI induction reaching 3.0-fold at 120 min after SI induction (**Figure 2B**). In plastids, significant increases in roGFP2-Orp1 oxidation state were detected from 5 minutes after SI induction onwards increasing by 2.2-fold 1 hour after the SI induction (**Figure 2C**). This demonstrates that SI stimulates increased H_2_O_2_ oxidation in the cytosol, mitochondria and plastids. SI did not stimulate roGFP2-Orp1 oxidation in all organelles; no significant alterations in R_405/488_ were measured in peroxisomes or nuclei (**Figure 2D-E**). To obtain further information on early increases in roGFP2-Orp1 in the cytosol, mitochondria and plastids following the SI induction, we tracked multiple individual pollen tubes over a 20-minute period. While control pollen tubes treated with growth medium maintained a steady R_405/488_, the kinetics of R_405/488_ increases in SI-induced pollen tubes varied, but all increased within 10 minutes after induction (**Figure 2F-H**). In untreated growing pollen tubes, representative time-lapse images showed that in the cytosol, roGFP2-Orp1 was more reduced in the tip region than in the shank and no major changes in oxidation state were observed during growth (**Figure 3A-C, Video 1**). After SI induction, this tip-to-shank difference rapidly disappeared, with a relatively uniform increase in roGFP2-Orp1 oxidation observed throughout the pollen tube (**Figure 3D-F, Video 2**). This oxidation occurred shortly after the inhibition of tip growth (**Figure 3D**, **Video 2)**. SI induction also triggered a general increase in roGFP2-Orp1 oxidation in mitochondria and plastids, although no distinct spatial features were observed in these organelles in either growing or SI-induced pollen tubes (Supplementary **Figure S3**).

**Figure 2.**
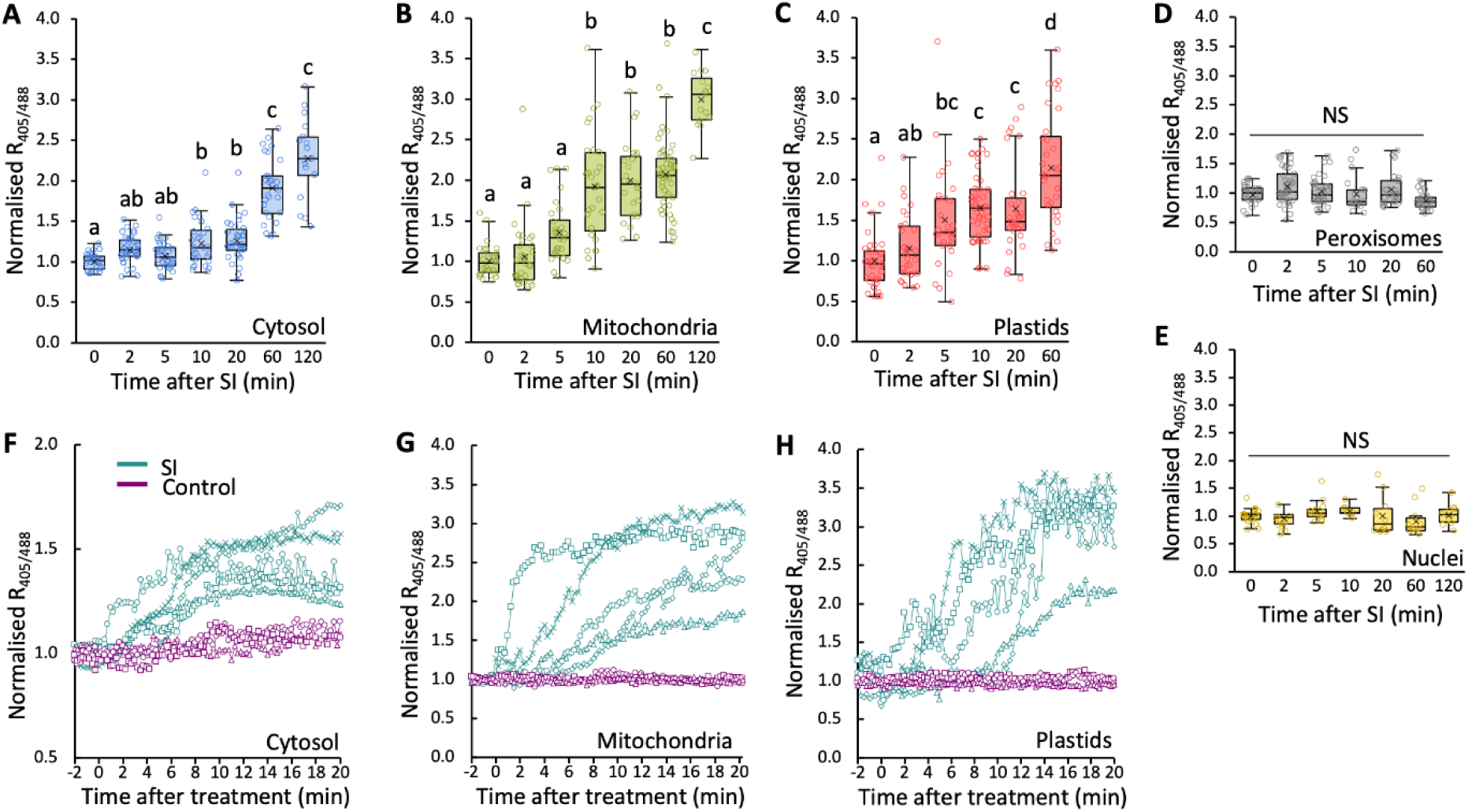
Changes in oxidation state of roGFP2-Orp1 targeted to various subcellular compartments in Arabidopsis pollen tubes. (A-E) Quantification of R_405/488_ of roGFP2-Orp1 targeted to the cytosol (A), mitochondria (B), plastids (C), peroxisomes (D) and nuclei (E) after SI induction. (A-C) R_405/488_ increased significantly (P < 0.001, one-way ANOVA) in the cytosol (A), mitochondria (B) and plastids (C) after SI induction compared with untreated (0 min) samples. (D, E) No significant increase (NS, *P* > 0.05, one-way ANOVA) in R_405/488_ was detected in peroxisomes (D) or nuclei (E) after SI induction compared with untreated (0 min) samples. (F-H) Quantification of R_405/488_ of roGFP2-Orp1 targeted to the cytosol (F), mitochondria (G) and plastids (H) during 0-20 min after SI induction (SI; n = 5) or treatment with growth medium (control; n = 3) in representative Arabidopsis pollen tubes expressing PrpS_1_ and roGFP2-Orp1. All R_405/488_ measurements were normalised to the mean ratio of untreated samples. For A-E, roGFP2-Orp1 was stabilised using 20 mM NEM at the end of each reaction. Different letters indicate significant differences based on Tukey’s test. Sample sizes: (A): n = 16-42; (B) n = 15-51; (C) n = 29-49; (D) n = 19-43; (E) n = 8-21.

**Figure 3.**
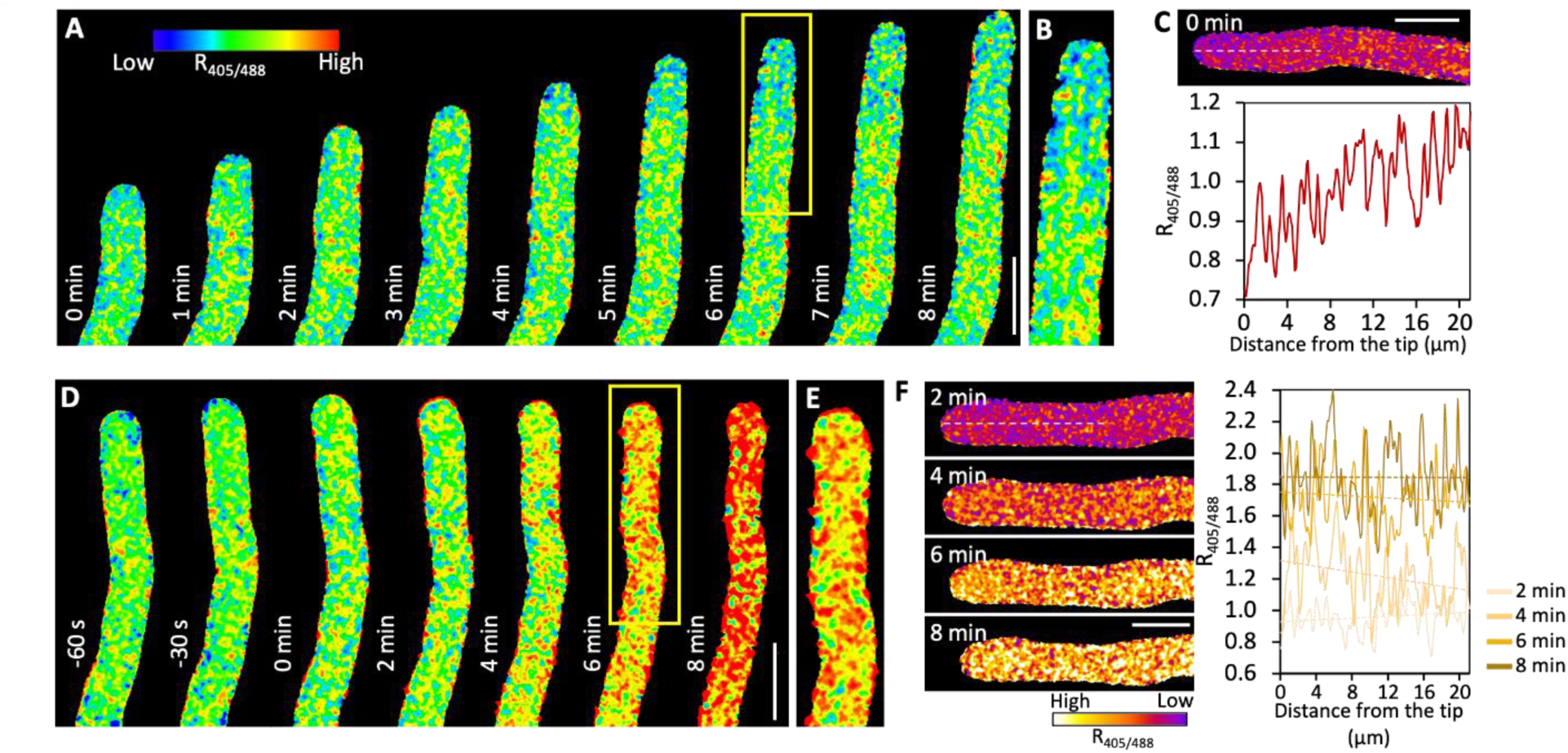
Increases in cytosolic roGFP2-Orp1 oxidation state in Arabidopsis pollen tubes during the SI response. (A, D) Representative ratio images of Arabidopsis pollen tubes expressing PrpS_1_ and roGFP2-Orp1, either untreated (A, UT) or treated with PrsS_1_ to induce SI (D, SI). Scale bar = 10 μm. (B, E) Enlarged views of the regions indicated by yellow boxes in (A) and (D), showing the spatial variation in roGFP2-Orp1 oxidation before (B) and after SI induction (E). (C, F) Fluorescence ratio profiles of cytosolic roGFP2-Orp1 (R_405/488_) along the white dashed lines in a representative pollen tube without treatment (C) and after the SI induction (F). The profiles illustrate SI induced dynamic changes in roGFP2-Orp1 oxidation state over time. Scale bar = 10 μm.

To examine whether the SI-induced oxidation of roGFP2-Orp1 was solely due to increased H_2_O_2_ production or also involved impaired reduction capacity of the sensor due to other SI-induced event, we measured roGFP2-Orp1 oxidation following treatment with 2.5 mM H_2_O_2_, with or without a preceding SI induction. No significant differences in roGFP2-Orp1 oxidation were observed in the cytosol or mitochondria after H_2_O_2_ treatment, regardless of SI induction (Supplementary **Figure S4A-B**). However, in the plastids, roGFP2-Orp1 was significantly more oxidised when H_2_O_2_ treatment followed SI induction, compared to H_2_O_2_ treatment alone (Supplementary **Figure S4C**). These results suggest that the increases in roGFP2-Orp1 oxidation in the cytosol and mitochondria after SI induction are due to elevated H_2_O_2_ production. In contrast, the rise in roGFP2-Orp1 oxidation in plastids might be partly due to a compromised plastid reduction capacity following SI induction.

### The electron transport chain contributes to SI-induced H_2_O_2_ elevation in mitochondria

Mitochondrial ROS is generated as a by-product of the electron transport chain (ETC) during respiration. To investigate the contribution of the ETC to H_2_O_2_ production during SI in the pollen tube, we used antimycin A, an inhibitor of complex III, to disrupt the ETC during SI responses (Wagner et al., 2019; Rattanawong et al., 2021). Treatment of growing pollen tubes with antimycin A for 20 minutes resulted in ATP depletion (**Figure 4A**) but did not significantly affect roGFP2-Orp1 oxidation state in the cytosol and mitochondria of growing pollen tubes (**Figure 4B-C**). Importantly, antimycin A mitigated the SI-induced increases in roGFP2-Orp1 oxidation in mitochondria and the cytosol after 20 min of SI induction (**Figure 4B, C**). This supports the hypothesis that the ETC contributes to SI-induced increases in H_2_O_2_ production in mitochondria and that this mitochondrial H_2_O_2_ can diffuse into the cytosol. A similar trend was observed for plastids, where SI-induced oxidation was partially (although not significantly) reduced by antimycin A (**Figure 4D**). Consistent with ETC inhibition and mitochondrial dysfunction, antimycin A also caused a rapid decrease in the mitochondrial membrane potential within 5 min of treatment (Supplementary **Figure S5**).

**Figure 4.**
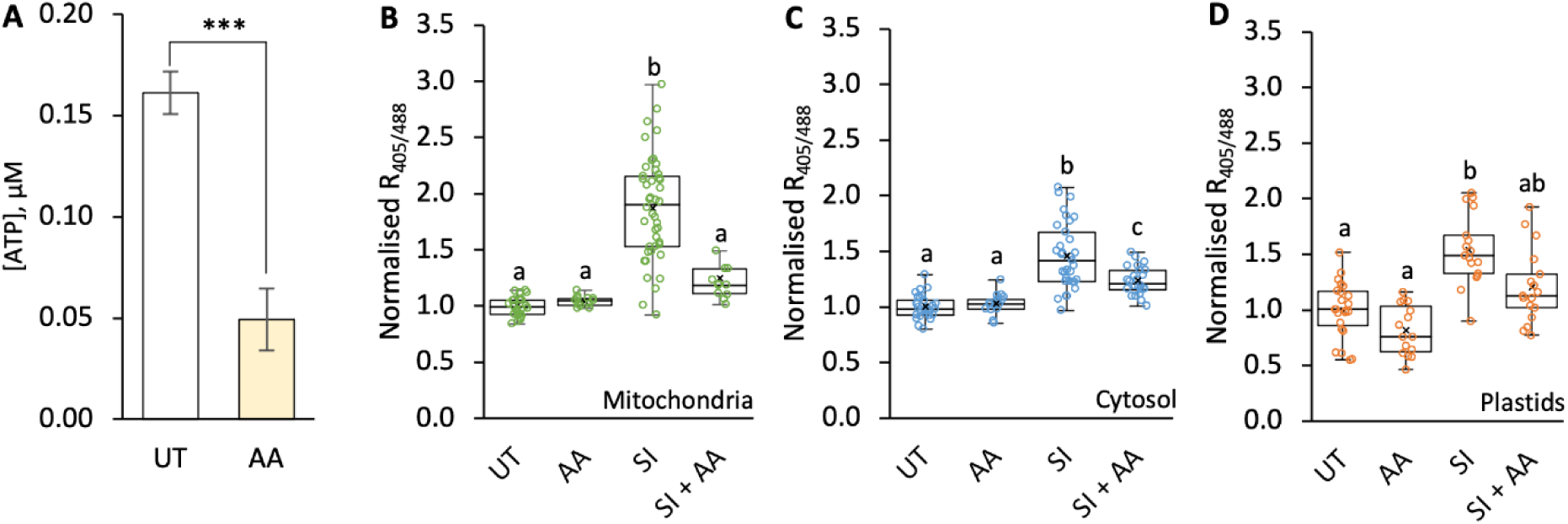
Inhibition of the electron transport chain (ETC) mitigates SI-induced ROS elevation in Arabidopsis pollen tubes. (A) Quantification of ATP levels in untreated (UT) pollen tubes (empty bars) or 20 minutes after the addition of 10 μM antimycin A (AA, filled bars). ATP levels were significantly reduced following treatment (\*\*\**P* < 0.001, one-way ANOVA; n = 3). (B-D) Quantification of 405/488 nm fluorescence ratio (R_405/488_) of roGFP2-Orp1 targeted to mitochondria (B), cytosol (C) and plastids (D) in untreated (UT) pollen tubes or 20 minutes after treatment with AA, with or without SI induction. n ≥ 13. Different letters indicate significant (*P* < 0.001) differences based on Tukey’s test.

### SI and H_2_O_2_ induce the collapse of mitochondrial membrane potential (Δψ_m_)

Our data demonstrating the contribution of the ETC to SI-induced mitochondrial H_2_O_2_ production suggest that ETC enzyme activities of complex I to IV remain (at least partially) active within the first 20 min after SI induction. However, we previously showed that SI induces ATP depletion within just a few minutes (Wang et al., 2022). This indicates that the ETC becomes uncoupled from ATP synthesis very early during the SI response. To investigate this, we treated pollen tubes with H_2_O_2_ and measured rapid decreases in cellular ATP levels, falling to 37.8% of the original level within 2 min, and to 10.3% by 15 min (**Figure 5A**). Thus, a possible explanation of ETC-ATP synthesis uncoupling is that SI-induced H_2_O_2_ causes a collapse of Δψ_m_, which is required by the ATP synthase (complex V) to generate ATP.

**Figure 5.**
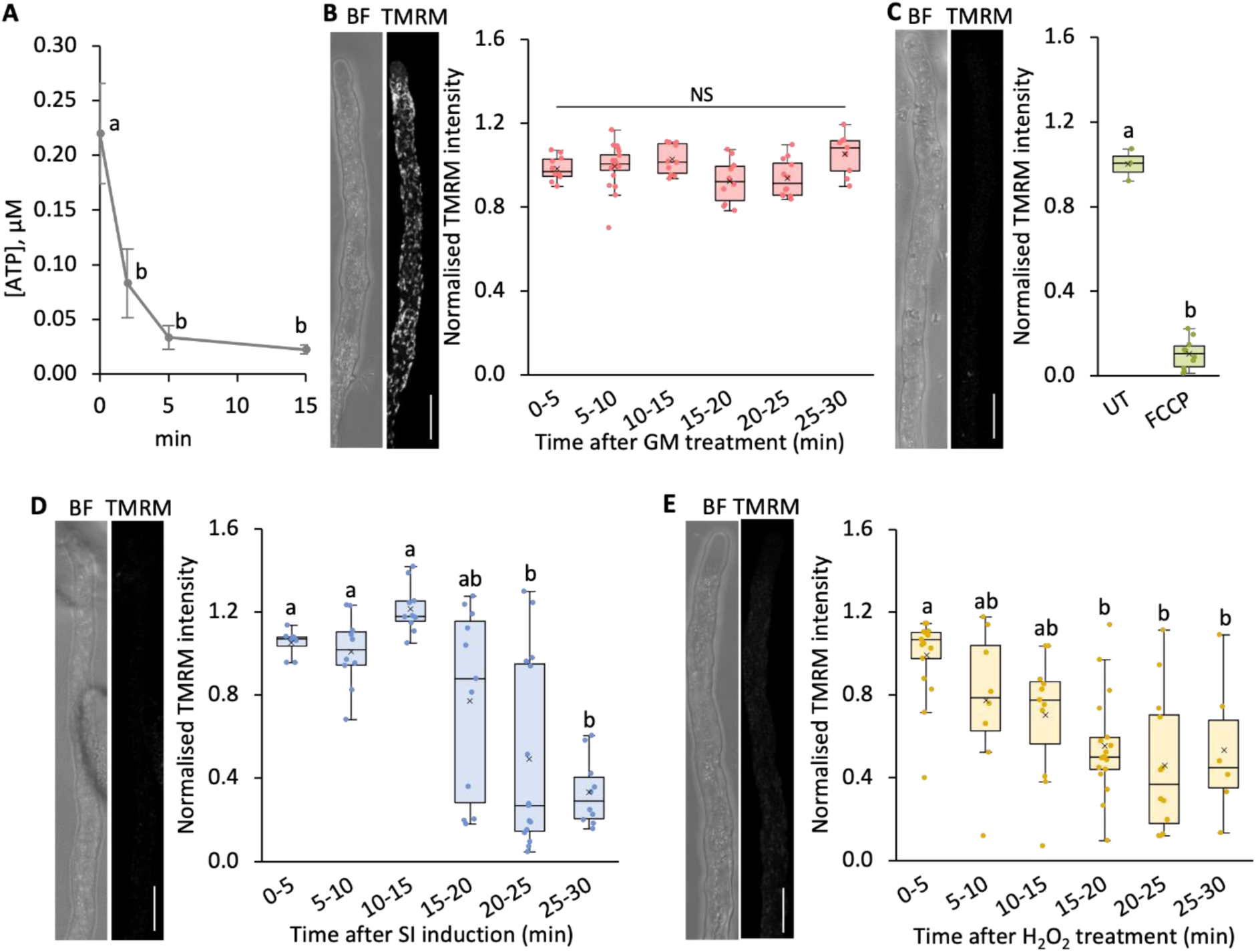
SI and H_2_O_2_ induce decreases in mitochondrial membrane potential (Δψ_m_) in Arabidopsis pollen tubes. (A) Quantification of ATP levels in pollen tubes treated with 2.5 mM H_2_O_2_. (B-E) Left panels show representative bright field (BF) and TMRM fluorescence images of pollen tubes after 20 minutes (B–D) or 10 minutes (E) of treatment. Scale bar = 10 μm. Right panels show the corresponding quantification of TMRM fluorescence in mitochondria after treatment with growth medium (GM) (B), FCCP (C), SI induction (D) and treatment with 2.5 mM H_2_O_2_ (E). Different letters indicate significant differences based on Tukey’s test (A and C, *P* < 0.001; D, *P* < 0.05; E, *P* < 0.01). Samples sizes: A, n =3; B, n = 9-20; C, n = 3-10; D, n = 8-13; E, n = 6-17.

To measure Δψ_m_ in pollen tubes, we used the well-established Δψ_m_ probe, tetramethyl rhodamine methyl ester (TMRM) (Perry et al., 2011) along with carbonyl cyanide-p-trifluoromethoxyphenylhydrazone (FCCP), a potent uncoupler of Δψ_m_ (Khan et al., 2024). In growing pollen tubes, a bright, stable TMRM signal was detected, indicating a healthy mitochondrial Δψm (**Figure 5B**). Treatment with FCCP for 10 min resulted in a significantly diminished TMRM signal (**Figure 5C**), confirming the probe’s responsiveness to changes in Δψ_m_. After SI induction, TMRM fluorescence was significantly reduced within 15-20 min, to 77.2 ± 45.0 % of the initial level, with a further decrease to 33.1 ± 16.2 % by 25-30 min (**Figure 5D**), demonstrating collapse of Δψ_m_. Similarly, treatment with 2.5 mM H_2_O_2_ caused TMRM fluorescence to decrease to 55.3% by 15-20 min, remaining relatively stable thereafter (**Figure 5E**). This demonstrates that both SI and H_2_O_2_ induce collapse of Δψ_m_.

### Manipulation of [Ca^2+^]_cyt_ alters SI-induced collapse of Δψ_m_ and roGFP2-Orp1 oxidation

[Ca^2+^]_cyt_ dynamics are important for mitochondrial calcium homeostasis and influence Δψ_m_ (Duchen et al., 2000; Loro et al., 2012). To investigate the relationship between SI-induced increases in [Ca^2+^]_cyt_ and collapse of Δψ_m_, we simultaneously imaged [Ca^2+^]_cyt_ and Δψ_m_ in pollen tubes using the genetically encoded [Ca^2+^]_cyt_ indicator YC3.6 and the TMRM probe. Tracking individual pollen tubes revealed that increases in YC3.6 R_venus/CFP_ ratios occurred before decreases in TMRM fluorescence (**Figure 6A**). This suggests that the SI-induced [Ca^2+^]_cyt_ elevation is an upstream event leading to the collapse of Δψ_m_. To further explore this link, we compared the effects of SI induction and calcium ionophore (A23187) treatment, in combination with the calcium channel blocker GdCl_3_. Both SI and A23187 treatments induced substantial increases in [Ca^2+^]_cyt_ (**Figure 6C**; Supplementary **Figure S6**) and led to substantial decreases in TMRM fluorescence intensity after 25-40 min compared to untreated pollen tubes (**Figure 6B**). However, GdCl_3_ maintained TMRM fluorescence intensity in SI-induced pollen tubes at the same level as untreated pollen tubes (105.7 ± 8.0 %, *P* = 0.23; **Figure 6B-C**), confirming that Ca^2+^ influx is required for the SI-induced collapse in Δψ_m_.

**Figure 6.**
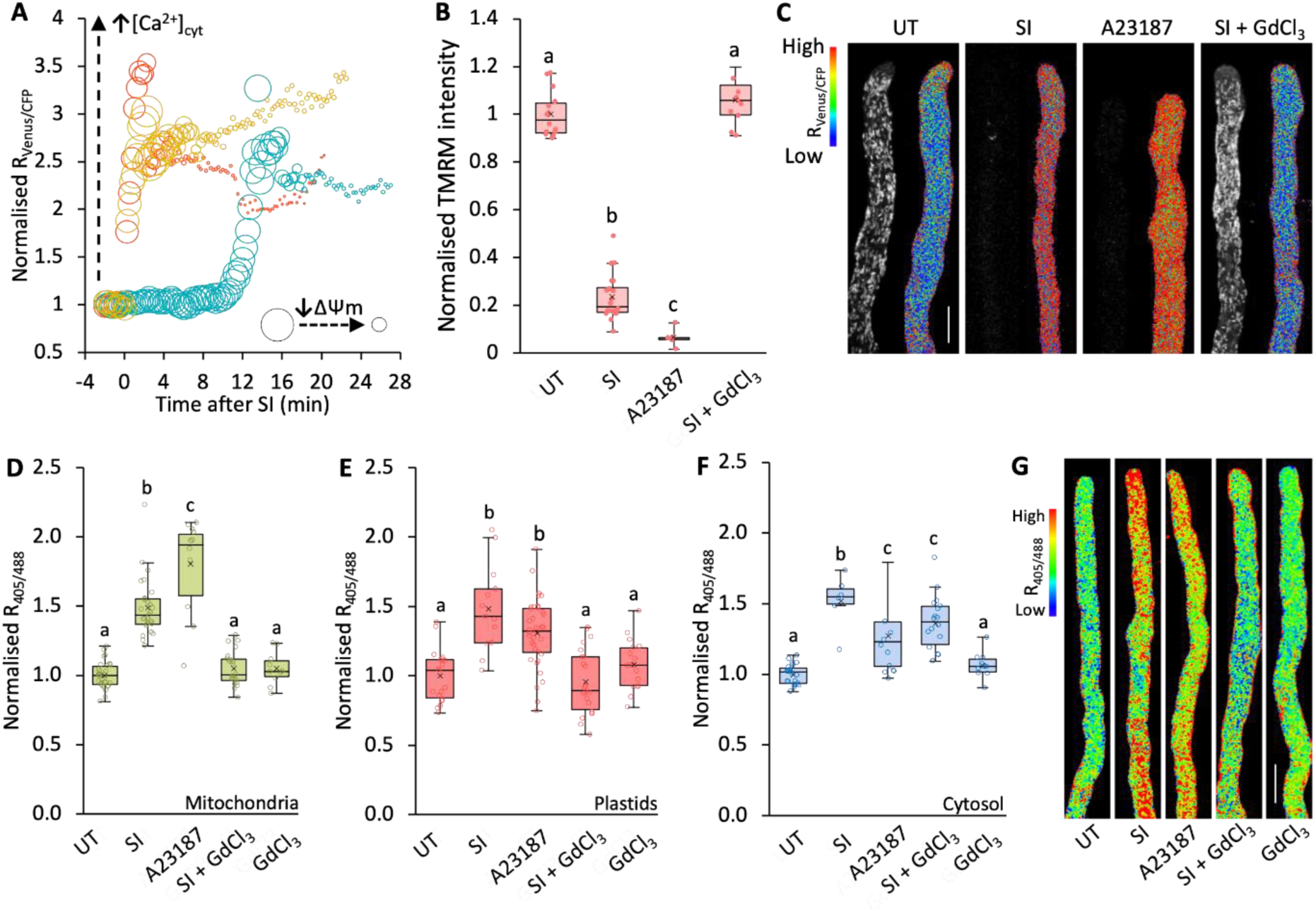
SI-induced roGFP2-Orp1 oxidation and mitochondrial ΔΨ_m_ are dependent on increases in [Ca^2+^]_cyt_. (A) Quantification of YC3.6 fluorescence ratio (R_Venus/CFP_) and TMRM fluorescence intensity in three representative Arabidopsis pollen tubes expressing PrpS_1_ and YC3.6 following treatment with PrsS_1_ (SI induction). A higher R_Venus/CFP_ indicates increased [Ca^2+^]_cyt_. Smaller bubble widths indicate lower TMRM fluorescence intensity, which reflects a decrease in ΔΨ_m_. Data from individual pollen tubes are shown in gold, red, and teal. (B) Quantification of TMRM fluorescence intensity in mitochondria in untreated (UT) pollen tubes and at 25-40 min after treatment with PrsS_1_ (SI), A23187, or PrsS_1_ plus GdCl_3_. n = 14, 20, 6, 22, respectively. (C) Representative images show YC3.6 fluorescence ratio (R_Venus/CFP_; right panels) and TMRM fluorescence signal (left panels) in untreated (UT) pollen tubes and 20 minutes after treatment with PrsS_1_ (SI), A23187 or PrsS_1_ plus GdCl_3_. Scale Bar, 10 μm. (D-F) Quantification of 405/488 nm fluorescence ratio (R_405/488_) of roGFP2-Orp1 targeted to mitochondria (D), plastids (E), and cytosol (F) in untreated (UT) pollen tubes or 20 minutes after treatment with PrsS_1_ (SI), A23187, PrsS_1_ plus GdCl_3_, or GdCl_3_ only. Sample sizes: D, n = 10-30; E, n = 13-36; F, n = 9-22, respectively). (G) Representative ratio images of pollen tubes expressing PrpS_1_ and roGFP2-Orp1 under the same conditions as in (F). Scale bar, 10 μm. In B, D, E and F, different letters indicate significant (*P* < 0.05) differences based on Tukey’s test.

To examine possible links between [Ca^2+^]_cyt_ and ROS in mitochondria, plastids and the cytosol, we quantified roGFP2-Orp1 oxidation state in PrpS_1_ expressing pollen tubes treated with A23187 and GdCl_3_. In mitochondria and plastids, both SI induction and A23187 treatment significantly increased roGFP2-Orp1 oxidation (**Figure 6D-E**). SI induction in the presence of GdCl_3_ prevented the SI-induced roGFP2-Orp1 oxidation in both mitochondria and plastids; the R_405/488_ ratios in these compartments were not significantly different to untreated pollen tubes (*P* = 0.84 for mitochondria and *P* = 0.98 for plastids; **Figure 6D-E**). In the cytosol, the effect of GdCl_3_ on SI-induced roGFP2-Orp1 oxidation was less pronounced than in the mitochondria and plastids (**Figure 6F**). Pollen tubes displayed elevated roGFP2-Orp1 oxidation (R_405/488_) at 20 minutes after SI induction (1.67 ± 0.28-fold, *P* < 0.001) or A23187 treatment (1.27 ± 0.25, *P* < 0.001). Co-treatment with GdCl_3_ significantly impaired SI-induced roGFP2-Orp1 oxidation compared to SI alone (*P* < 0.001; **Figure 6F-G**). Together these data show that Ca^2+^ influx and increases in [Ca^2+^]_cyt_ are required to achieve elevation of SI-induced ROS across these compartments, implicating Ca^2+^ signalling upstream of ROS increases.

### Manipulation of cytosolic pH modulates SI-induced decrease in Δψ_m_

Since SI also triggers cytosolic acidification (Wilkins et al., 2015; Wang et al., 2022), we investigated whether changes in [pH]_cyt_ affect alterations in the oxidation state of roGFP2-Orp1 in the cytosol and mitochondria. Artificially reducing [pH]_cyt_ to 5.5 using propionic acid significantly increased roGFP2-Orp1 oxidation (R_405/488_) in both the cytosol and mitochondria compared to untreated pollen tubes (**Figure 7A**). Conversely, clamping the [pH]_cyt_ at 7 with propionic acid prevented SI-induced roGFP2-Orp1 oxidation in both the cytosol and mitochondria compared to untreated pollen tubes (**Figure 7B**). These treatments also affected Δψ_m_ (**Figure 7C**). SI induction and lowering [pH]_cyt_ to 5.5 decreased TMRM fluorescence intensity. In contrast, clamping [pH]_cyt_ at 7 prevented SI-induced decreases in TMRM fluorescence. These findings firmly implicate that cytosolic acidification plays a crucial role in the signalling network underlying these processes. To assess whether H_2_O_2_ participates in a feedback loop affecting [pH]_cyt_, we treated pollen tubes with 2.5 mM H_2_O_2_, which decreased [pH]_cyt_ to 6.69 compared to untreated pollen tubes with a [pH]_cyt_ of 7.28 (**Figure 7D**), though the acidification was less pronounced than that induced by SI (Wang et al., 2022). Together these results indicate that cytosolic acidification contributes to SI-induced mitochondrial dysfunction and H_2_O_2_ generation.

**Figure 7.**
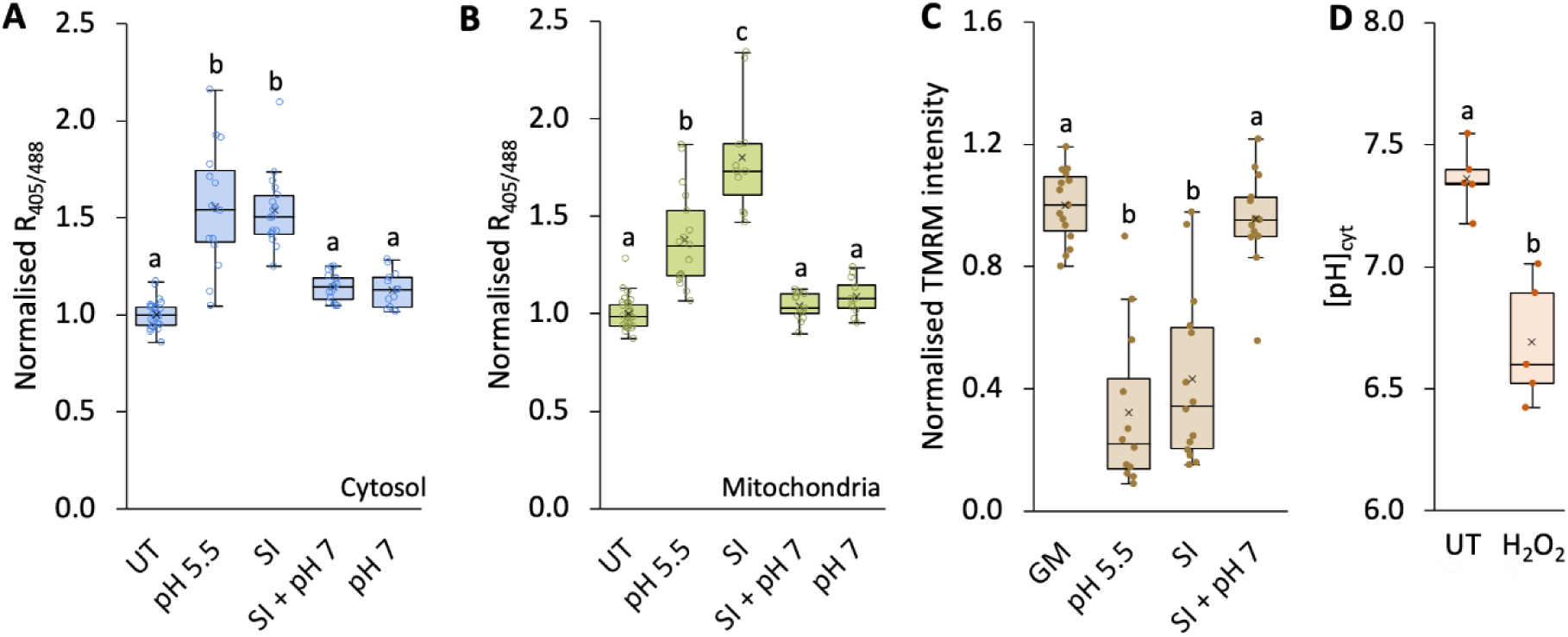
Manipulation of cytosolic pH modulates SI-induced mitochondrial dysfunction in Arabidopsis pollen tubes. (A, B) Quantification of 405/488 nm fluorescence ratio (R_405/488_) of roGFP2-Orp1 targeted to the cytosol (A) and mitochondria (B) in untreated (UT) pollen tubes and at 20 minutes following treatment with propionic acid at pH 5.5, PrsS_1_ (SI), PrsS_1_ with propionic acid at pH 7 (SI + pH 7), or propionic acid at pH 7 alone. n ≥ 11. (C) Quantification of TMRM fluorescence intensity in mitochondria of pollen tubes treated with growth medium (GM), PrsS_1_ (SI), PrsS_1_ with propionic acid at pH 7 (SI + pH 7), or propionic acid at pH 5.5 for 25 minutes. Treatment at pH 5.5 triggered decreases in mitochondrial ΔΨ_m_ similar to those induced by SI while clamping at pH 7.0 prevented SI-induced decreases in mitochondrial ΔΨ_m_. n ≥ 12. (D) Quantification of cytosolic pH ([pH]_cyt_) in pollen tubes 25 minutes following treatment with 2.5 mM H_2_O_2_, which induced cytosolic acidification. n = 5. Different letters in A-D indicate significant (*P* < 0.001) differences based on Tukey’s test.

### SI and H_2_O_2_ inhibit respiratory metabolism in pollen tubes

We previously identified irreversible oxidation of several enzymes associated with glycolysis, organic acid metabolism and mitochondrial ATP synthase that were induced by both SI and H_2_O_2_ (Haque et al., 2020). As SI triggers rapid ATP depletion, this suggested significant energetic and metabolic changes occur in incompatible pollen tubes (Wang et al., 2022). Our data indicate that SI triggers increased mitochondrial H_2_O_2_ generation and decreased mitochondrial membrane potential. To explore the impact of SI on respiratory metabolism we measured oxygen uptake, the activity of the glycolytic enzyme glyceraldehyde 3-phosphate dehydrogenase (GAPDH) and the concentration of TCA cycle intermediates in response to SI and H_2_O_2_.

To assess whether SI and ROS inhibit respiration, we measured the oxygen consumption of *Papaver* pollen tubes using an oxygen electrode. Using *Papaver* pollen tubes was necessary, due to the limited amount of Arabidopsis pollen available. Following SI induction, oxygen uptake rates significantly decreased, dropping to 59.7% at 30-60 min and to 30% at 60-90 min compared to growth medium treated controls (**Figure 8A**). Following treatment with 2.5 mM and 10 mM H_2_O_2_, oxygen uptake rates decreased to 71.6% and 59.2%, respectively, during the 30-60 min period compared to untreated samples (**Figure 8B**).

**Figure 8.**
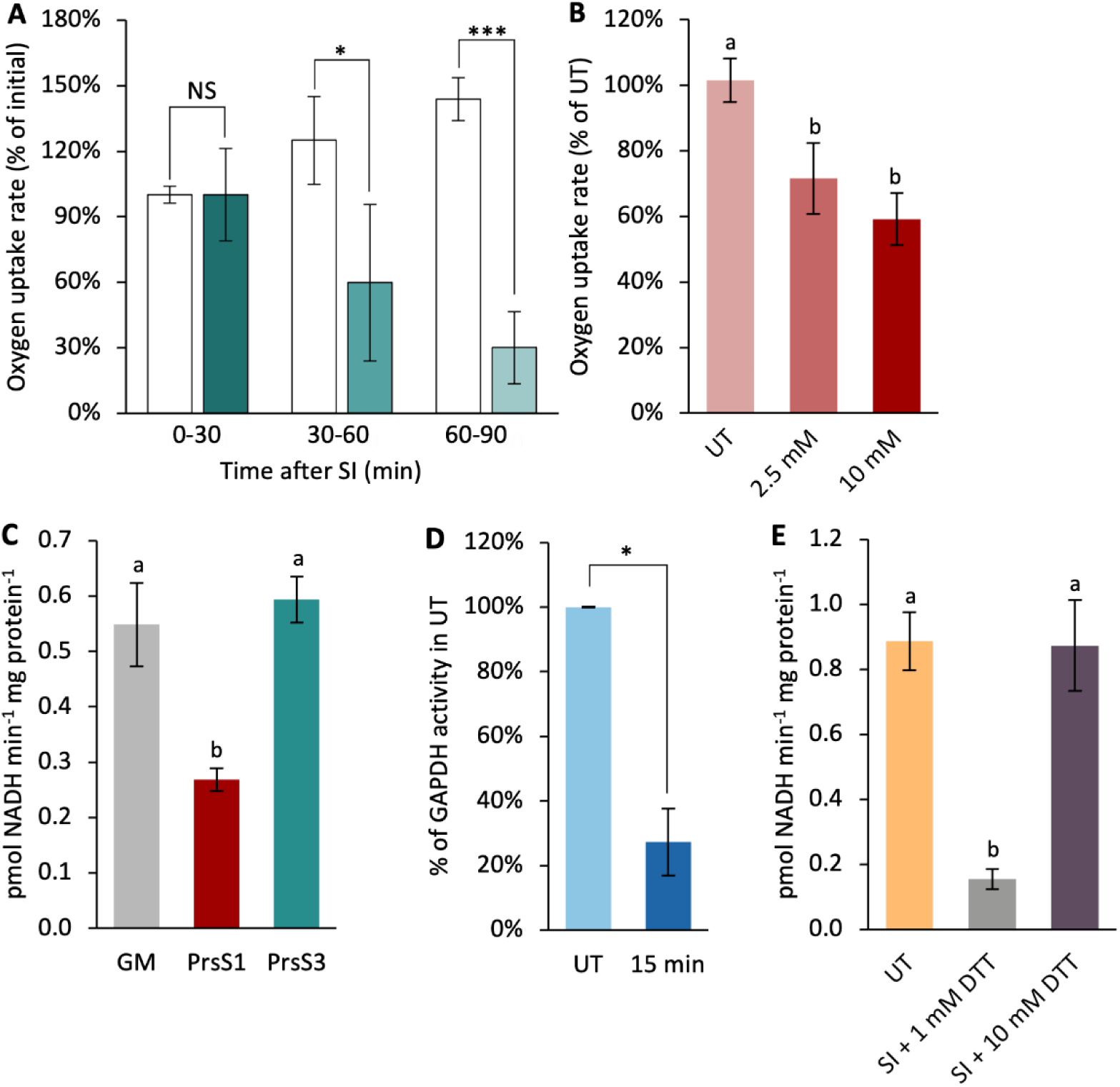
SI and H_2_O_2_ inhibit glycolysis and mitochondrial respiration in pollen tubes. (A) Oxygen uptake rates in *Papaver* pollen tubes during three time intervals: 0-30 min, 30-60 min, and 60-90 min after treatment with growth medium (control, empty bars) or after SI induction (filled bars). Rates were normalised to mean values during the 0-30 min interval. NS, not significant. \**P* < 0.05, \*\*\**P* < 0.001, one-way ANOVA. n = 4. (B) Oxygen uptake rates in untreated (UT) *Papaver* pollen tubes and those treated with 2.5 mM or 10 mM H_2_O_2_ for 30 minutes. Rates were normalised to mean values of untreated samples. (n = 4). (C) GAPDH activities in Arabidopsis pollen tubes measured 5 min after treatment with growth medium (GM), PrsS_1_ (SI induction) or PrsS_3_ (n = 3). (D) GAPDH activities in Arabidopsis pollen tubes 15 minutes after treatment with 2.5 mM H_2_O_2_. \**P* < 0.05, Mann-Whitney U test (n = 4). (E) DTT prevented SI-induced inactivation of GAPDH in Arabidopsis pollen tubes. n ≥ 3. B, C and E, Different letters indicate significant differences based on Tukey’s test (B, *P* < 0.01, C and E, *P* < 0.001).

As GAPDH was previously identified as a target for oxidative post-translational modifications in *Papaver* pollen tubes triggered by both SI and H_2_O_2_ (Haque et al., 2020), we measured its activity in Arabidopsis pollen extracts. 5 min after SI induction, GAPDH activity was significantly reduced to 48.9% of the activity in control samples. This effect was SI-specific as a compatible treatment had no significant effect on GAPDH activity relative to control (GM) samples (**Figure 8C**). This demonstrates that SI induces rapid impairment of GAPDH activity in incompatible pollen tubes. Treatment of pollen tubes with H_2_O_2_ also led to a decrease in GAPDH activity. Within 15 min of treatment, GAPDH activity was reduced to 27% of the level in untreated pollen tubes (**Figure 8D**). As addition of 10 mM dithiothreitol (DTT) restored GADPH activity (**Figure 8E**), this suggests that SI and H_2_O_2_ inactivate GAPDH by oxidation of cysteine thiol groups to sulfenic acids, while further oxidation of the sulfenic acid gives rise to the previously reported oxidative post-translational modifications (Haque et al., 2020). Decreased GAPDH activity following SI induction or H_2_O_2_ treatment was also detected in *Papaver* pollen tubes (Supplementary **Figure S7**). These findings suggest that SI-induced inactivation of GAPDH is mediated by elevated levels of ROS stimulated by SI. Further, glutathione (GSH), a prominent component of the thiol antioxidant system, decreased within 15 minutes of SI (**Figure 9A**), supporting the occurrence of wider redox perturbation. This suggests that glycolysis and the supply of pyruvate to the TCA cycle is rapidly decreased during SI. Consistent with this conclusion, the concentration of 2-oxoglutarate, succinate and malate had decreased by ∼50% within 15 minutes of initiating SI with PrsS_1_ but not in the compatible response with PrsS_3_ (**Figure 9B-D**). In a separate experiment, 2-oxoglutarate decreased within 15 mins, while malate and succinate showed a progressive (but not significant) decrease from 15 to 120 min post SI induction (Supplementary **Figure S8**). Intriguingly, the TCA cycle acids at the beginning of the cycle (citrate/isocitrate and aconitate) were unaffected (**Figure 9E-F** and Supplementary **Figure S8C**). These findings demonstrate that both SI and elevated H_2_O_2_ disrupt key metabolic pathways, rapidly impairing energy production and redox homeostasis in pollen tubes.

**Figure 9.**
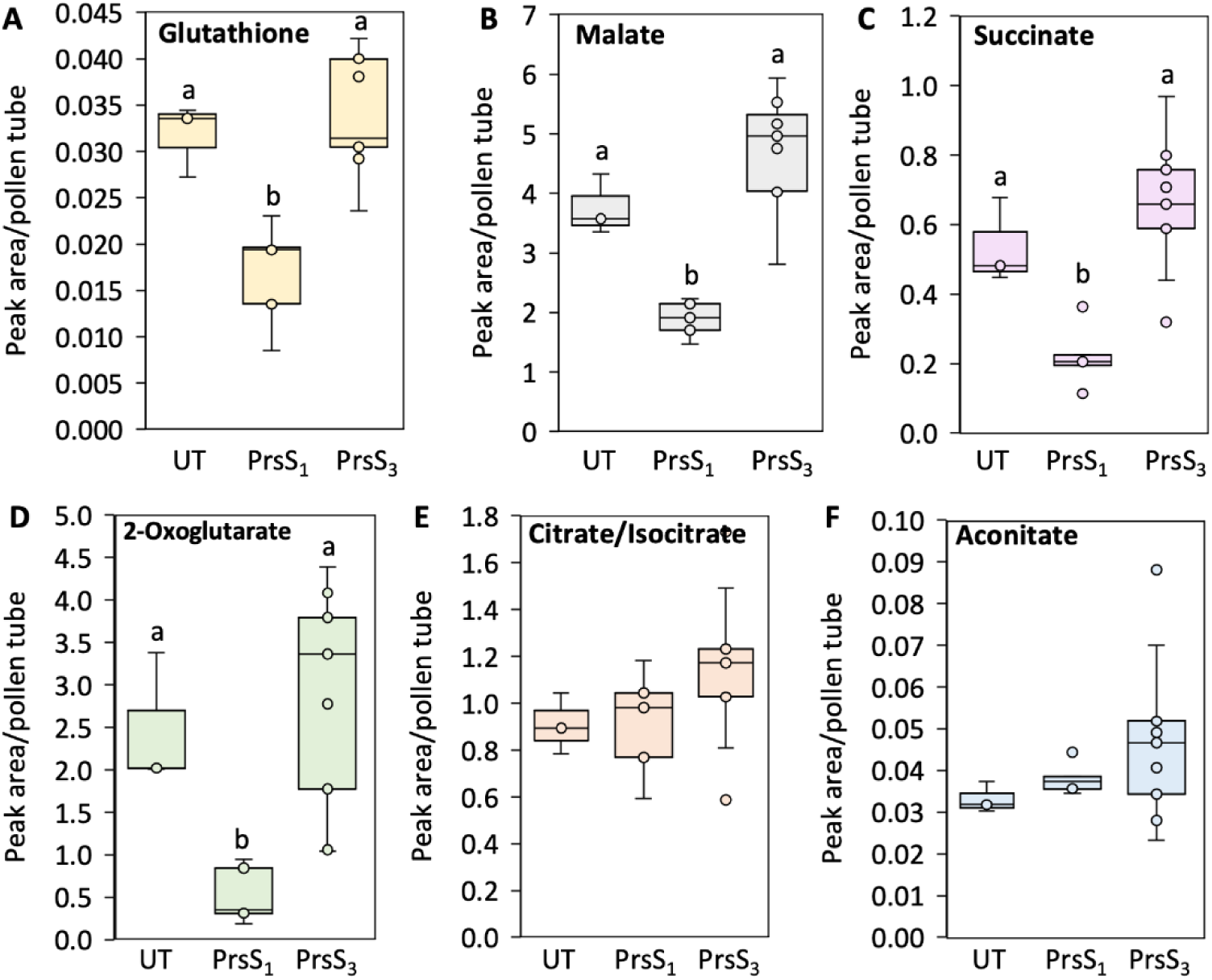
SI induces decreases in some TCA cycle acids and glutathione in Arabidopsis pollen tubes. Levels of glutathione (A), malate (B), succinate (C) and 2-oxoglutarate (D) were significantly reduced in pollen tubes 15 minutes after SI induction. In contrast, levels of citrate/isocitrate (E) and aconitate (F) remained unchanged. Sample sizes: Untreated (UT), n = 3; PrsS_1_, n = 5; PrsS_3_, n= 9. Different letters indicate significant (*P* < 0.001) differences based on Tukey’s test.

## Discussion

Using the heterologous Arabidopsis system, which exhibits key features of *Papaver* SI (Lin et al., 2020; Wang et al., 2020), we provide an integrated, subcellular-resolution analysis of the early SI response in pollen. This has revealed that the SI response disrupts mitochondrial electron transport, triggers compartment-specific H₂O₂ accumulation and suppresses glycolytic flux *via* oxidation-dependent inactivation of GAPDH.

Previously, we showed that SI in *Papaver* stimulates rapid ROS production and H₂O₂-dependent protein oxidation (Wilkins et al., 2011; Haque et al., 2020). Here, using the genetically encoded H_2_O_2_ biosensor roGFP2-Orp1 (Gutscher et al., 2009; Nietzel et al., 2019; Ugalde et al., 2021; Arnaud et al., 2023), we dissected H_2_O_2_ dynamics across five subcellular compartments, identifying mitochondria as the primary site of SI-induced ROS generation and the likely origin of H_2_O_2_ increases in the cytosol and plastids. Importantly, we also show that ROS required for pollen tube growth, produced *via* NADPH oxidase activity at the tip, is functionally and spatially distinct from the SI-induced ROS. This provides one of the first clear demonstrations of dual ROS systems with opposing roles operating within the same plant cell type (the vegetative cell of the pollen tube). This multi-layered disruption occurs within 20 minutes of SI induction and is orchestrated through Ca^2+^ influx and a rapid rise in [Ca^2+^]_cyt_ followed soon after by cytosolic acidification, mitochondrial membrane depolarisation, ATP depletion, and redox perturbation within the pollen tube. Pharmacological and biosensor data suggest that mitochondrial dysfunction is both a source and a target of redox and pH perturbations, setting up a feedback loop that accelerates cellular shutdown. While previous work has identified individual SI-induced events, our study unites these into a mechanistic framework that links early signal transduction initiated by SI to rapid metabolic failure and oxidative stress within the first 10-20 minutes, which is well before PCD execution.

### SI-induced ROS is distinct from NADPH oxidase-sourced ROS necessary for tip growth

The NADPH oxidase isoforms RBOHH and RBOHJ, localised at the plasma membrane of the pollen tube tip, are required for pollen tube growth in Arabidopsis. Pollen tubes of mutants lacking both isoforms (*rbohH/J*) are prone to bursting and fail to reach the female gametophyte (Boisson-Dernier et al., 2013; Kaya et al., 2014). NADPH oxidase forms superoxide in the apoplast, which is rapidly dismutated to H_2_O_2_. Apoplastic NADPH oxidase-dependent superoxide production at the pollen tube tip has previously been detected by NBT reduction in tobacco (Potocký et al., 2007), and we found the same pattern in both *Papaver* and Arabidopsis. As anticipated, initiation of SI caused a very rapid loss of tip-localised superoxide production in both systems, coinciding with pollen tube growth arrest. NADPH oxidase activity is subject to complex control *via* Ca^2+^, phosphoinositides and Rho-related GTPases from plants (ROPs) (Potocký et al., 2003; Potocký et al., 2012). NADPH oxidase was inhibited by the Ca^2+^ channel blocker Gd^3+^, confirming a role for Ca^2+^ influx in its activation. The rapid SI-induced inactivation of NADPH oxidase is most likely due to disruption of the normal [Ca^2+^]_cyt_ signature of tip-high, oscillating [Ca^2+^]_cyt_ that is required for growth (Franklin-Tong et al., 1997). Our results demonstrate that the ROS required for normal pollen tube tip growth, originating from NADPH oxidase activity, are functionally distinct from the mitochondrially produced ROS induced by SI. This is consistent with studies showing that cytosolic H_2_O_2_ levels are only marginally affected by NADPH oxidase activity (Boisson-Dernier et al., 2013; Arnaud et al., 2023). This represents one of the first clear examples of two spatially distinct ROS sources playing divergent functional roles within a single cell type, the vegetative cell of the pollen tube, which drives tip growth and contains the generative and vegetative nuclei.

### Mitochondria are the source of the SI-induced H_2_O_2_ production

To track intracellular H_2_O_2_ dynamics during the SI response, we used the roGFP2-Orp1 biosensor targeted to various organelles of Arabidopsis pollen co-expressing PrpS. The biosensor was functional, responding as expected to both H_2_O_2_ and the reductant dithiothreitol. Importantly, roGFP2-Orp1 remains functional at pH 5.5 (Nietzel et al., 2019), so the measured oxidation state is unaffected by the cytosolic acidification induced by SI or treatment with propionic acid. Consistent with observations in leaves (Arnaud et al., 2023), peroxisomes exhibited the most oxidised redox state, followed by intermediate oxidation in mitochondria and plastids, and the most reduced state in the cytosol and nucleus. Initiation of SI caused oxidation of the biosensor in the cytosol, mitochondria, and plastids, but not in the nuclei and peroxisomes. Since roGFP2-Orp1 reports the redox balance between oxidation by H_2_O_2_ and its reduction by the thiol system, the signal is to some extent a reflection of both processes (Nietzel et al., 2019). The kinetics of oxidation were very similar in each compartment, with oxidation becoming evident after ∼10 minutes. This is similar to the time course of CM-H_2_DCF oxidation previously observed after triggering SI in *Papaver* (Wilkins et al., 2011), but the use of roGFP-Orp1 has enabled the subcellular location of oxidation to be established. Since H_2_O_2_ can diffuse across membranes, facilitated by aquaporins (Bienert and Chaumont, 2014; Rodrigues et al., 2017), its source is not immediately obvious. However, the mitochondrial electron transport inhibitor antimycin A, which inhibits complex III, prevented SI-induced roGFP2-Orp1 oxidation in mitochondria and cytosol. This suggests that SI-induced H_2_O_2_ production is dependent on mitochondrial electron transport. Although it is reported that antimycin A can increase ROS production, the effect is not always large (Maxwell et al., 1999; Moller, 2001) nor consistent. For example, antimycin A treatment of Arabidopsis seedlings did not result in rapid oxidation of cytosolic roGFP-Orp1 (Khan et al., 2024), but in pollen, SI-induced oxidation is rapid and is inhibited by antimycin A across mitochondria and cytosol. Thus, our results strongly suggest that mitochondria are the primary source of SI-induced H_2_O_2_, likely *via* residual electron transport, with ROS subsequently diffusing into the cytosol and plastids. It is also possible that the decreased cytosolic pH inhibits H_2_O_2_ scavenging enzymes such as ascorbate peroxidase and could contribute to SI-induced H_2_O_2_ accumulation. In particular, the plastid form of ascorbate peroxidase has a steep decrease of activity below its pH 8.0 optimum (Mittler and Zilinskas, 1991).

Overall, the results are consistent with SI causing a disruption in mitochondrial metabolism that results in oxygen reduction to superoxide and consequent H_2_O_2_ production. H_2_O_2_ rapidly moves to the cytosol and plastids. The rapid oxidation observed in plastids is unlikely to result from ROS generation within the organelle itself, since plastids do not have an electron transport chain (Renato et al., 2015). Instead, plastid ROS accumulation most likely reflects diffusion of H_2_O_2_ from mitochondria, potentially *via* aquaporins. As plastids support rapid pollen tube growth *via* plastid glycolysis, the oxidative pentose phosphate pathway, and fatty acid synthesis (Emes and Neuhaus, 1997), their redox state may be additionally affected by SI-induced growth arrest. The lack of a response in peroxisomes may reflect their specialisation in housing H_2_O_2_-producing oxidases along with high catalase activity (Smirnoff and Arnaud, 2019).

### SI decreases glycolysis by oxidative inactivation of GAPDH and inhibits the TCA cycle

SI and exogenous H_2_O_2_ caused a decrease in oxygen uptake by *Papaver* pollen, demonstrating that SI has a large impact on respiration. Several lines of evidence point to disrupted metabolism, including decreased ATP, reduced GAPDH activity (affecting glycolysis in both cytosol and plastids), a decrease in some TCA cycle intermediates, and a decrease in the mitochondrial membrane potential. We previously identified redox-sensitive post-translational modifications in two glycolytic enzymes, GAPDH and enolase, following both SI and H_2_O_2_ treatment in *Papaver* pollen (Haque et al., 2020), implicating oxidative damage to these proteins. Here we show that some of these modifications have functional significance since we found that NAD-GAPDH activity is rapidly decreased during SI and following H_2_O_2_ treatment. Crucially, this inactivation was reversed by DTT, indicating that reversible cysteine oxidation is involved. GADPH is well known to be under redox control, undergoing reversible oxidation by H_2_O_2_ on a conserved cysteine residue (Cys155), resulting in inactivation (Schneider et al., 2018; Dumont and Rivoal, 2019). As well as influencing glycolysis rate in the cytosol and plastids, the oxidised form of GADPH can translocate to the nucleus and may be involved redox signalling, while the reduced form associates with voltage-dependent anion channels (VDAC) on the outer mitochondrial membrane, possibly as part of a glycolytic complex (Dumont and Rivoal, 2019). The oxidative inactivation of glycolysis will impose a limitation on ATP generation by substrate level phosphorylation and decrease supply of pyruvate to the mitochondria. The latter will limit TCA cycle activity, as shown by the decrease in TCA cycle intermediates beyond citrate/isocitrate. Moreover, given that SI triggers a decrease in glycolysis, the possibility of ATP generation by fermentation as suggested by Wang et al. (2022) is unlikely. This conclusion is reinforced for Arabidopsis, whose pollen has a low fermentation capacity (Liu et al., 2022). In support of an early impact on metabolism, previous work in *Papaver* has shown that soluble inorganic pyrophosphatases (sPPases), which are essential for biosynthetic reactions, become phosphorylated and inactivated within minutes of SI induction (Rudd et al., 1996; de Graaf et al., 2006), further illustrating the rapid shutdown of key metabolic processes following SI. Together, our data establish a direct mechanistic link between SI-induced ROS and impaired metabolism *via* redox regulation of GAPDH and suggest that oxidative modifications of other glycolytic enzymes may also contribute. While some studies, such as (Obermeyer et al., 2013), have inferred changes in pollen metabolism in response to mitochondrial dysfunction using proteomic data, to date direct evidence for ROS-mediated inactivation remains scarce in plants. These findings therefore provide important insights into how redox signalling can modulate energy metabolism in response to cellular stress signals such as SI.

### SI triggers multi-layered cellular disruption

SI induction very rapidly triggers H_2_O_2_ production and disruption to respiratory metabolism. The previously described increases in [Ca^2+^]_cyt_, cytosolic acidification and decrease in ATP levels are equally rapid, making it difficult to unravel cause and effect (Wilkins et al., 2011; Wilkins et al., 2015; Wang et al., 2019; Haque et al., 2020; Wang et al., 2022;). However, integrating our findings enables us to propose a model outlining the likely sequence and interdependence of these early events during the first 10-20 minutes of the SI response (**Figure 10**). Elevated [Ca^2+^]_cyt_ is the earliest detectable change (Bosch and Franklin-Tong, 2024). Pharmacological manipulation using Ca^2+^ channel blockers and ionophores indicate that several key events, including decrease in Δψ_m_, increased roGFP2-Orp1 oxidation (reflecting H_2_O_2_ production), decreased ATP levels, and cytosolic acidification, are downstream of the initial influx and subsequent increases in [Ca^2+^]_cyt_. Furthermore, manipulation of [pH]_cyt_ shows that acidification increases H_2_O_2_ and decreases Δψ_m_ while these SI-induced events are blocked by maintaining neutral [pH]_cyt_. Acidification may therefore act as a direct and early effector in the SI response. To our knowledge, this represents one of the first demonstrations in plants that cytosolic acidification can directly impair mitochondrial function. While the importance of pH homeostasis for mitochondrial activity is well recognized, direct evidence linking cytosolic acidification to mitochondrial dysfunction is a physiological context has been lacking. We propose/hypothesize that SI triggers a combined disruption of Δψ_m_, most likely *via* inhibition of mitochondrial electron transport, alongside oxidative inhibition of glycolysis and TCA cycle. Together, these effects would lead to decreased respiration and ATP production. Notably, a small release of cytochrome c in the cytosol was detected within 10 minutes after SI induction in Papaver pollen tubes, preceding the more pronounced increase observed after one hour (Thomas and Franklin-Tong, 2004). This early release could contribute to a decrease in electron transport rate. The resulting drop in ATP may, in turn, potentiate cytosolic acidification, at least in part *via* decreased plasma membrane and/or vacuolar H^+^-ATPase activity. However, this alone is unlikely to account for the dramatic acidification observed following SI, suggesting that addition mechanisms must be involved (Wang et al., 2022). A recent characterisation of the pollen mitochondrial proteome (Boussardon et al., 2025) has revealed unique features associated with high respiration rates needed for rapid pollen tube tip growth including high abundance of TCA cycle enzymes, electron transport complexes and Ca^2+^ homeostasis and a deficiency of transcriptional and translational machinery. These specialised features may render pollen highly susceptible to the kind of disruption observed during SI. Collectively, these findings support a model in which mitochondrial dysfunction and metabolic inhibition form a self-reinforcing feedback loop, amplifying cellular disruption.

**Figure 10.**
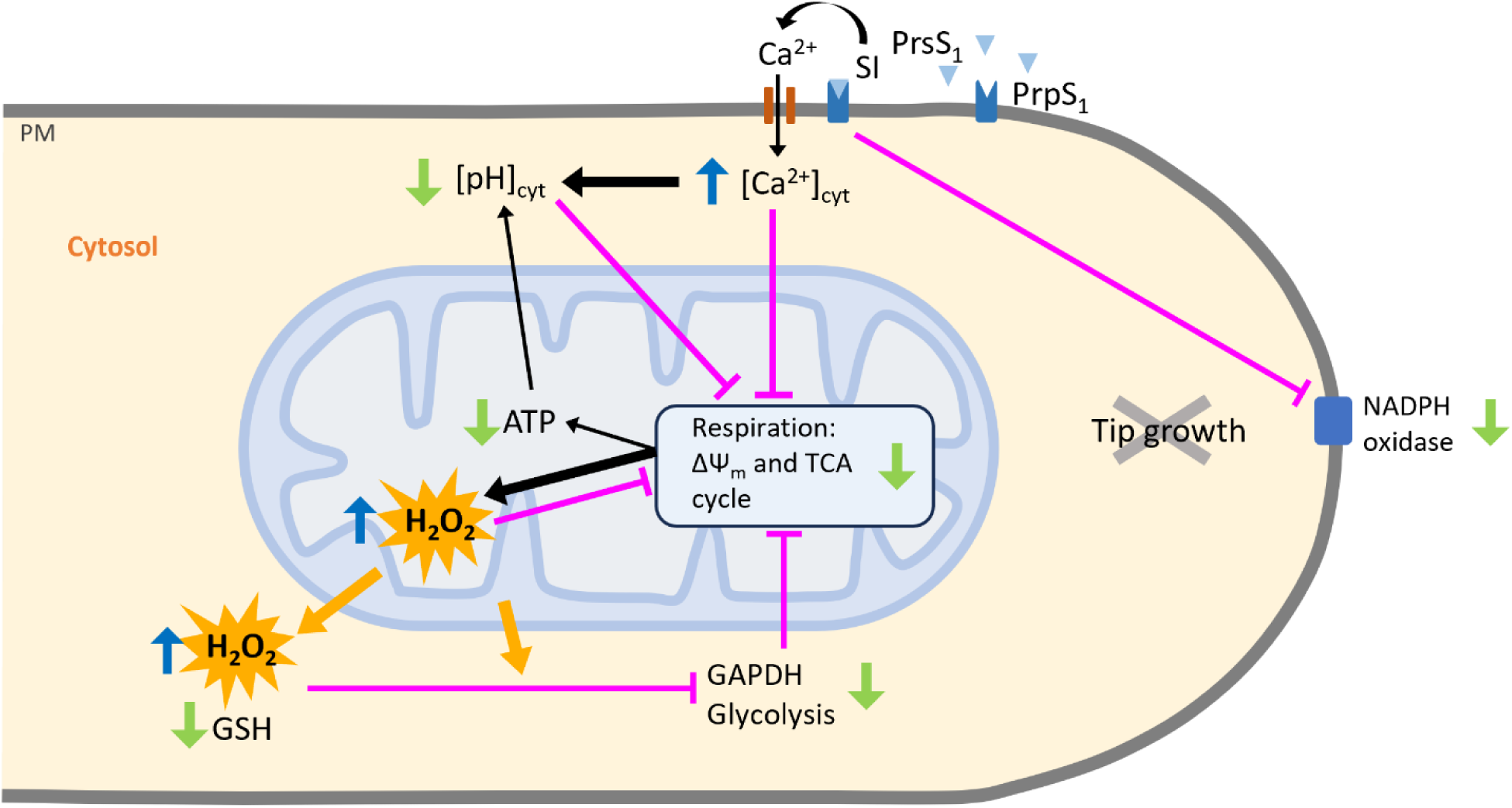
A summary of the interconnected events occurring during the first ten to twenty minutes of the SI response in pollen tubes. We previously showed that SI is initiated when the stigmatic *S*-determinant PrsS (PrsS_1_ in the diagram) binds to its cognate pollen plasma membrane protein PrpS_1_. This triggers a rapid influx of Ca²⁺, resulting in elevated cytosolic Ca²⁺ concentration ([Ca²⁺]_cyt_), disruption of the tip-focused Ca²⁺ gradient and arrest of pollen tube tip growth. Downstream of the [Ca²⁺]_cyt_ increase, a network of intracellular responses is rapidly triggered (within 10-20 min). including cytosolic acidification and decreased ATP concentration.

### Linking early mitochondrial disruption to PCD

SI triggers PCD downstream of early signals (Thomas and Franklin-Tong, 2004; Wang et al., 2019; Bosch and Franklin-Tong, 2024). Although the current study did not directly examine PCD, several of the rapid events triggered by SI, such as mitochondrial dysfunction, ROS production, and ATP depletion, have previously been associated with PCD in plants (Xie et al., 2014; Van Aken and Van Breusegem, 2015; Huang et al., 2016). While the role of mitochondria in plant PCD remains rather contentious and an area of ongoing debate, it has been suggested that they can act as early integrators of PCD signalling in certain systems (Curtis and Wolpert, 2004; Mur et al., 2008; Moller et al., 2021). Our previous studies on *Papaver* have shown that SI induces the release of cytochrome *c* within 10 minutes, followed by dramatic morphological changes, including swelling and loss of cristae (Geitmann, et al., 2004; Thomas and Franklin-Tong, 2004). These alterations resemble mitochondrial changes associated with apoptosis in animal cells, and similar mitochondrial morphology transitions have also been linked to PCD in plants (Scott and Logan, 2008; Ocampo-Hernández et al., 2022). Combined with our findings of ATP depletion and mitochondrial dysfunction, this demonstrates that mitochondria are early targets of SI-induced signalling to PCD.

Studies to date suggest that many plant PCD processes, such as those in the stigma, tapetum or hypersensitive response, involve vacuolar rupture and NADPH oxidase-derived ROS (Franklin-Tong and Bosch, 2021; Zhang et al., 2021; Kacprzyk et al., 2024). However, our findings here demonstrate that mitochondrially-derived ROS contributes to metabolic dysfunction that underpins PCD in the SI system. This, together with previous data, clearly implicates very early mitochondrial alterations observed within minutes (Thomas and Franklin-Tong, 2004) as being involved as a causal event upstream of PCD (and not just a marker of dying cells). Consistent with this, recent work in Arabidopsis root cap cells showed that mitochondrial disruption, including the release of matrix proteins, is one of the earliest detectable events during developmental cell death, preceding the breakdown of other organelles (Wang et al., 2024). Thus, there are clearly distinct signals and outputs that are responsible for PCD, underlining the diversity of plant PCD systems. Establishing how rapid changes in intracellular Ca²⁺, pH, and redox state influence the downstream commitment to PCD remains a key challenge for the field.

### Evidence for ROS signatures

We have provided a clear demonstration that spatially and temporally distinct ROS (and Ca^2+^) signals can serve as intracellular signatures that allow for functionally divergent roles within the pollen tube, essentially a single cell type, resulting in distinct cellular responses and cell fates: tip growth or SI. While the concept and evidence for distinct spatial-temporal Ca^2+^ “signatures” are well-established, very few ROS signatures within single cell types have been identified in plants. It is known that different ROS sources can give rise to different responses. Examples include NADPH oxidase-derived superoxide/H_2_O_2_ in pathogen responses and tip growth of root hairs and pollen (Torres et al., 2002; Foreman et al., 2003; Potocký et al., 2007; Arnaud et al., 2023), singlet oxygen and hydrogen peroxide (*via* chlorophyll excitation and electron transport respectively) in chloroplast to nucleus retrograde signalling (Exposito-Rodriguez et al., 2017; Page et al., 2017) and H_2_O_2_ production from mitochondrial electron transport in retrograde signalling and PCD (Van Aken et al., 2015; Khan et al., 2024). An interaction between NADPH oxidase and mitochondrial-derived ROS has been implicated in stomatal closure (Postiglione and Muday, 2023) and the PAMP response in Arabidopsis involves oxidation *via* NADPH oxidase-dependent and independent ROS sources (Arnaud et al., 2023). Together, these findings support a model in which the timing, localisation, and source of ROS generation are critical in determining specific physiological outcomes, even within a single cell.

## Conclusions

We have provided the first integrated analysis of the complex cascade of cellular and metabolic events that involve potential positive feedback loops triggered during the early stages of the SI response (**Figure 10**), using a combination of cell imaging and metabolic assays. Focusing on the first 20 minutes of this response, we show that a Ca^2+^-and pH-dependent series of events results in mitochondrial ROS production, likely *via* disruption of mitochondrial electron transport. This results in inhibition of mitochondrial respiration and ATP production. Concurrently, oxidative signals are observed in the cytosol, oxidising and inhibiting the glycolytic enzyme GAPDH, further impacting ATP production. Our data implicate the mitochondria as a central hub of SI-triggered metabolic collapse and ROS signalling that is triggered by increases in [Ca^2+^]_cyt_ and [pH]_cyt_. However, the precise molecular mechanisms by which the rapid Ca^2+^ and pH changes perturb mitochondrial function and ROS production remain to be elucidated, representing an important area for future investigation.

## Methods

### Plant material and growth conditions

Transgenic *Arabidopsis thaliana* lines expressing PrpS_1_ in pollen, along with various fluorescent markers (see Supplementary Table S1) were used in this study. Plants were grown in controlled environment chambers under a 16-hour light/8-hour dark photoperiod, with a light intensity of 135 μmol m⁻² s⁻¹. Mature pollen grains were harvested from fully opened flowers. *Papaver rhoeas* pollen grains were collected from field-grown plants and stored over silica gel at -20 °C before use, as described previously (Snowman et al., 2002).

### Plasmid construction and generation of transgenic plants

Details of primers, targeting sequences for roGFP2-Orp1 and Golden Gate modules used are listed in Supplementary Table S2.

The dual-expression vector carrying *ProNTP303:PrpS_1_-ProNTP303:LifeAct-mRuby2* was generated using GreenGate cloning (Lampropoulos et al., 2013). Promoter *NTP303* was amplified using primer sets *F-A-NTP303/R-B-NTP303*, *F-D-NTP303/R-E-NTP303* with the vector carrying *ProNTP303:PrpS_1_-GFP* as the template (de Graaf et al., 2012). The PCR products were cloned into pJET1.2 using CloneJET PCR Cloning Kit (ThermoFisher) to obtain the entry vectors *pEN-A-ProNTP303-B* and *pEN-D-ProNTP303-E*. Generation of other entry vectors, including *pEN-B-PrpS1-C*, *pEN-C-tRBCS-D*, *pEN-E-LifeAct-mRuby2-F* and *pEN-F-tMAS-G,* was described in (Lin et al., 2020). These entry clones were assembled into GreenGate destination vector *pFAST-GK-AG* to produce the dual-expression vector carrying *ProNTP303:PrpS_1_-ProNTP303:LifeAct-mRuby2*.

Constructs for roGFP2-Orp1 localisation in the cytosol, mitochondria, plastids, peroxisomes and nuclei were assembled using Golden Gate reactions (Weber et al., 2011; Engler et al., 2014), following the procedures described in Arnaud et al. (2023) with the following modifications.

To ensure pollen-specific expression, the CaMV 35S promoter (*Pro35S*) was replaced with the pollen-specific promoter *ProNTP303* (Weterings et al., 1995). *ProNTP303* was amplified using primer sets NTP303_F/NTP303_AATG_R and NTP303_F/NTP303_TAACT_R as described above. These amplified fragments were cloned into pICH41233 and pICH41295, respectively. The pICH41295-ProNTP303 module was used for assembling constructs containing nuclear localisation signal (NLS), whereas the pICH41233-ProNTP303 module was used for assembling constructs containing a mitochondria-targeting signal (MTS), chloroplast (Chl)- and peroxisome-targeting signal serine-lysine-leucine (SKL).

The introduction of targeting sequences for the localisation of roGFP2-Orp1 to subcellular compartments followed procedures as described in Arnaud et al., 2023. Except for the MTS, a sequence encoding the first 54 amino acids of the transit peptide of the Arabidopsis homologue (At5g08680) of the *Nicotiana plumbaginifolia* ATP synthase beta-3 subunit (with 98% sequence identity, Chaumont et al., 1994; Carrie et al., 2015) was amplified using Arabidopsis Col-0 cDNA as the template with primers MTS-_F and MTS-_R. The chloroplast-targeting signal was amplified from pICH78133 (Engler et al., 2014) using primers Chloro_F and Chloro_R. These fragments were subsequently cloned into pICH41246.

The plasmid carrying *ProNTP303:Lifeact-mRuby2-ProNTP303:PrpS_1_* was transformed into *Arabidopsis thaliana* Col-0. Constructs for roGFP2-Orp1 localisation were transformed into Arabidopsis plants carrying *ProNTP303:Lifeact-mRuby2-ProNTP303:PrpS_1_*. *Agrobacterium tumefaciens* GV3101 and the floral dip method (Clough and Bent, 1998) were used for transformation. Transgenic seedlings were selected and screened based on fluorescence intensity for strong expression, which were used for further analyses.

### Pollen tube growth and treatment

*Arabidopsis thaliana* and *Papaver rhoeas* pollen were germinated *in vitro* in liquid growth medium for over 60 min before treatment and imaging, as previously described (Snowman et al., 2002; Wang et al., 2020). Self-incompatibility responses were induced by adding recombinant PrsS_1_ to *Arabidopsis* pollen tubes expressing PrpS_1_ or PrsS_1_ and PrsS_3_ to *Papaver* pollen tubes (*S_1_S_3_*) to a final concentration of 20 μg ml^-1^ as described previously (Wilkins et al., 2015; Wang et al., 2020). H_2_O_2_ (Sigma-Aldrich) or dithiothreitol (DTT, Thermo Fisher Scientific) were applied at concentrations indicated in the results. To evaluate the effect of SI on the reductive capacity of roGFP2-Orp1, SI was induced for 1.5 minutes, followed by a 5-minute treatment with 2.5 mM H₂O₂ before imaging. To evaluate mitochondrial membrane potential, *Arabidopsis* pollen tubes were treated with 1 μM tetramethylrhodamine methyl ester (TMRM, Thermo Fisher Scientific) and washed with growth medium before imaging (Scaduto and Grotyohann, 1999; Perry et al., 2011). Various compounds (Sigma-Aldrich), including Diphenyleneiodonium chloride (DPI), antimycin A (AA), carbonyl cyanide-p-trifluoromethoxyphenylhydrazone (FCCP) and calcium Ionophore A23187, were applied at concentrations detailed in the results section. The carrying solvents, DMSO (except ethanol for AA), at a final concentration of no greater than 0.1 % (v/v) in growth medium, had no effect on pollen tube growth. GdCl_3_ and caffeine were applied at 500 μM and 10 mM, respectively. To preserve the redox state of roGFP2-Orp1 after PrsS or drug treatment, N-ethylmaleimide (NEM, Sigma-Aldrich) was used at 20 mM, as described by (Wendt et al., 2022). Cytosolic pH was altered using 50 mM propionic acid at pH 7.0 or 5.5, as previously described (Wilkins et al., 2015).

### Visualisation of tip-localised superoxide production

Superoxide production at the pollen tube tip was detected by nitroblue tetrazolium (NBT, Sigma-Aldrich) reduction at 5 mg ml^-1^ for 15 minutes. Images were captured using a Leica DMi8 microscope (100× CS2 objective, NA 1.40) equipped with a Leica TCS SPE camera. Grey values within the cytosolic region 0-50 μm from the apex were quantified using Fiji software (Schindelin et al., 2012).

### Confocal imaging and image analyses

Fluorescence markers were imaged using a Leica SP8 confocal microscope (×100 CS2 objective, NA 1.40). Lifeact-mRuby2 decorating F-actin, and sensors for [Ca^2+^]_cyt_ (Yellow Cameleon 3.6, YC3.6) and [pH]_cyt_ (pHGFP) were imaged as previously described (Wang et al., 2020). roGFP2-Orp1 was sequentially excited at 405 nm and 488 nm, with emission collected at 505-535 nm. For roGFP2-Orp1 targeting cytosol, the pinhole was set to 5 airy units (Nietzel et al., 2019), and for subcellular compartments, it was set to 1 airy unit. TMRM was excited at 561 nm and emission was collected at 650-775 nm. For each pollen tube, 11 z-stack images were taken with 0.9 μm intervals. Fluorescent intensity ratios (R_405/488_) for roGFP2-Orp1 were processed and quantified using Fiji software (Schindelin et al., 2012). The 0-40 μm from the pollen tube tip was selected as the region of interest (ROI). Background noise was subtracted using ‘Subtract background’ with radius of 50 pixels). Ratios for YC3.6 (R_Venus/CFP_) and pHGFP (R_405/488_) were processed and quantified using Leica Application Suite X (LAS X). Additional image processing for figure preparation was conducted using Fiji software (Schindelin et al., 2012). Measurements were performed on individual pollen tubes, with three independent experiments conducted per treatment.

### ATP assays

Intracellular ATP levels in *Arabidopsis* pollen tubes were quantified using the Luminescent ATP Detection Assay Kit (Abcam), as previously described (Wang et al., 2022).

### Measurement of pollen tube oxygen consumption

REDFLASH fibre optic oxygen sensors (FireSting-O_2_; PyroScience; Aachen, Germany) were used to measure oxygen uptake (Bénit et al., 2017). After being hydrated in a humid chamber for 50 minutes, *Papaver* pollen was grown in 2 ml glass vials containing liquid *Papaver* GM and well-sealed by caps allowing for the probe insertion and the addition of treatments including 2.5-10 mM H_2_O_2_ and the mixture of PrsS_1_ and PrsS_3_ or PrsS_3_ and PrsS_8_ (50-100 mg ml^-1^). The decrease in oxygen concentration with time was measured using the Pyro Oxygen Logger software. Respiration rate was determined by calculating the rate of oxygen uptake over 15 min periods normalised to weight of pollen.

### GAPDH activity assays

Glyceraldehyde 3-phosphate dehydrogenase (GAPDH) activity in pollen extracts was assessed by measuring NADH formation in the presence of glyceraldehyde 3-phosphate and arsenate. *Papaver* pollen was grown on solidified *Papaver* GM for two hours before the addition of treatments including 2.5 mM H_2_O_2_ and mixture of recombinant PrsS_1_ and PrsS_3_ proteins (50-100 mg ml^-1^) for 5 to 30 minutes. Arabidopsis pollen was grown on solidified Arabidopsis GM for 4-5 hours before the addition of treatments including 2.5 mM H_2_O_2_ and recombinant PrsS_1_ or PrsS_3_ proteins (50-100 mg ml^-1^) for 5 minutes. Pollen tube samples were collected in liquid nitrogen and ground into enzyme extraction buffer containing 50 mM HEPES/KOH pH 7.5, 10% (v/v) glycerol, 0.25% (w/v) BSA, 0.1% (v/v) Triton X-100, 10 mM MgCl_2_, 1 mM EDTA, 1 mM EGTA, 1x protease inhibitor cocktail, 1 mM PMSF and 1 mM DTT (Gibon et al., 2004). After centrifugation, enzyme extraction in the supernatant was quantified by the standard Bradford assay (Bio-Rad). Glycolytic GAPDH activity was monitored spectrophotometric at 340 nm *via* the production of NADH by using a ClarioStar plate reader (BMG LABTECH, Aylesbury, UK). The oxidation of glyceraldehyde-3-phosphate to 1,3-bisphosphoglycerate was measured in an assay buffer containing 20 mM Tris-HCl pH 7.5, 1.25 mM EDTA, 15 mM sodium arsenate, 1.25 mM NAD as described in (Hillion et al., 2017). After mixing the enzyme extraction with the assay buffer, the reaction was started by addition of 0.25 mM glyceraldehyde-3-phosphate in a 96-well UVStar microplate (Greiner Bio-One). Sodium arsenate was used as a co-substrate to form unstable 1-arseno,3-phosphoglycerate, whose degradation allows a favourable equilibrium for measuring the rate of GAPDH activity in the glycolytic forward reaction. The GAPDH specific activity was calculated from the increase in absorbance per minute using the extinction coefficient of NADH (6.22 mM^-1^ cm^-1^) and protein concentration.

### Pollen metabolites

*Arabidopsis* pollen tubes were germinated in growth medium for 2 hours before treatment. Pollen grains from 75 flowers were used per sample. Numbers of pollen tubes were estimated by measuring grey values within images of pollen tube growth field. Pollen tubes were collected by centrifugation at 6,000 *g* for 5 min at 22 °C before flash freezing using liquid nitrogen. Organic acids and glutathione were measured by LC-triple quadrupole MS/MS following derivatisation of pollen extracts with O-benzylhydroxylamine (Walvekar et al., 2018). 20 µl methanol/actetonitrile/0.1 M formic acid in water (4:4:2) was added to the pelleted pollen in 2 ml plastic centrifuge tubes followed by freeze thawing (-70°C) and sonication for 20 minutes in an ice-cold water bath. Derivatisation reagent (100 µl) was added, and samples incubated for 60 min. 20 µl of 0-20 µM of a standard mixture and blanks without pollen were also derivatised. The derivatisation reagent was prepared by mixing equal volumes of 1 M O-benzylhydroxylamine hydrochloride dissolved in acetonitrile/water (3:2) and 1 M N-(3-dimethylaminopropyl)-N′-ethylcarbodiimide dissolved in water/pyridine/37% HCl (16:1.6:1, pH 5.0). Ethyl acetate (300 µl) was added to each sample followed by vortexing and centrifugation (1,400 *g*, 5 min 4°C). The upper ethyl acetate phase containing the O-benzylhydroxylamine derivatives was transferred to tapered glass autosampler vials and dried down under vacuum, followed by dissolving in 20 µl 50% methanol. The samples were analysed with an Agilent 6420B triple quadrupole mass spectrometer coupled to a 1200 series Rapid Resolution HPLC system (Agilent Technologies, Santa Clara, USA). Samples and standards (5 µl) were injected onto a Polaris 3 C18-A (4.6 x 100 mm, 3.0 µm particle size) column with a Polaris 5 C18-A (10 x 2 mm) guard column (Agilent Technologies, Santa Clara, USA) column at 25°C and eluted at 0.3 ml min^-1^ with solvent A (0.1% (v/v) formic acid and 10% (v/v) methanol in water) and solvent B (0.1 % (v/v) formic acid in methanol). The elution gradient was 10% B for 0-0.5 min, 95% B from 11-15 min and 10% B at 26 min. The triple quadrupole source conditions were as follows: gas temperature 350°C, drying gas flow rate 9 l min^-1^, nebulizer pressure 35 psig, and capillary voltage 4 kV. Compounds were detected by multiple reaction monitoring in positive ion mode and the fragmentor voltage, and collision energies along with precursor and product ion m/z for each compound are shown in Table S3. Peak areas for each compound were extracted with MassHunter Quantitative Analysis software (v10.2, Agilent Technologies, Santa Clara, USA) and normalised to pollen number.

## Supporting information

Video 1. Time-lapse image of roGFP2-Orp1 oxidation state in a normally growing pollen tube. See Figure 3A and 3D for details.

Video 2. Time-lapse image of roGFP2-Orp1 oxidation state in a pollen tube during SI. See Figure 3A and 3D for details.

## Funding

This research was supported by the following grants from the Biotechnology and Biological Sciences Research Council (BBSRC): BB/T00486X/1 to MB and VF-T, BB/T005424/1 andto NS.

## Acknowledgments

We thank Dr. Deborah L. Salmon and Dr. Dyan Ankrett from the Exeter Mass Spectrometry Facility, Nicola Wood for laboratory technical assistance and Keith Ansell for assisting with plant growth.

## Author Contributions

LW, AH, JMD, VF, NS, and MB designed the research. LW, AH, JC, AR, ZL, and DA performed the research. LW, AH and NS analyzed the data. LW, VF, NS and MB wrote the original draft, and all authors were involved in final editing and review.

In this study, we show that tip-localised NADPH oxidase is inactivated, resulting in the inhibition of apical ROS production. SI triggers increased roGFP2-Orp1 oxidation in the cytosol and mitochondria. Mitochondria appear to be the primary source of SI induced increases in H₂O₂, and H₂O₂ generated here subsequently diffuses into the cytosol, leading to the large increases in cytosolic H₂O₂ observed in incompatible pollen tubes. This demonstrates that spatially and temporally distinct ROS signals can serve as intracellular signatures that allow for functionally divergent roles within the pollen tube.

Mitochondrial membrane potential (ΔΨₘ) decreases, concomitant with decreased ATP concentration. As manipulation of [pH]_cyt_ shows that acidification is sufficient to decrease ΔΨₘ and elevate H₂O₂, and maintaining neutral cytosolic pH prevents these effects, this suggests that cytosolic acidification acts as a direct and early effector in this SI response.

SI also results in rapidly reduced levels of antioxidants such as glutathione (GSH), demonstrating that redox homeostasis is further compromised. In addition, SI triggers oxidative inactivation of glyceraldehyde 3-phosphate dehydrogenase (GAPDH), a glycolytic enzyme. This will limit glycolytic flux and reduce pyruvate availability for mitochondrial respiration, contributing to decreased tricarboxylic acid (TCA) cycle activity thereby reducing ATP production in incompatible pollen tubes further. These findings provide insights into how redox signalling can modulate energy metabolism.

Decreased ATP may itself exacerbate cytosolic acidification *via* inhibition of ATP-dependent H⁺-ATPases at the plasma membrane and/or vacuole. We propose a model whereby these early SI-induced events establish a positive feedback loop in which SI-induced Ca²⁺ and pH changes disrupt mitochondrial function, which in turn generates further ROS and suppresses energy metabolism, amplifying intracellular disruption.

*Colour and symbol key:* Blue structure, mitochondrion. Black arrows denote activation/increases, magenta flat-end arrows denote inhibition. Green downward arrows indicate decreases, blue upward arrows indicate increases. Yellow bursts represent H₂O₂ accumulation. Blue box represents core mitochondrial functions. *Abbreviations*: SI, self-incompatibility; ΔΨm, Mitochondrial membrane potential; TCA, tricarboxylic acid cycle; GSH, Glutathione; GAPDH, glyceraldehyde 3-phosphate dehydrogenase, PM, plasma membrane; [Ca²⁺]_cyt_, cytosolic calcium concentration; [pH]_cyt_, cytosolic pH

**Video 1.** Time-lapse ratio-image series of a normally growing pollen tube from an *Arabidopsis thaliana* line co-expressing PrpS_1_, roGFP2-Orp1 and Lifeact-mRuby2 showing the distribution of roGFP2-Orp1 oxidation during normal growth. Fluorescence ratio (R_405/488_) scale refers to Figure 3A and 3D.

**Video 2.** Time-lapse ratio-image series of a normally growing pollen tube from an *Arabidopsis thaliana* line co-expressing PrpS_1_, roGFP2-Orp1 and Lifeact-mRuby2 showing the distribution of roGFP2-Orp1 oxidation during the SI response. Fluorescence ratio (R_405/488_) scale refers to Figure 3A and 3D.

**Figure S1. Disruption of tip-localized NADPH oxidase activity in *Papaver* pollen tubes following SI induction.** (A) Nitroblue tetrazolium (NBT) staining was used to detect superoxide production at the tips of *Papaver* pollen tubes during the SI response. Representative images show an untreated (UT) pollen tube and a pollen tube treated with PsS_1_ (SI) for 15 min. NBT staining was significantly reduced after SI induction. Scale bar = 10 μm. (B) Quantification of NBT staining intensity at pollen tube tips. Reduced NBT staining reflects decreased pixel intensity, indicating a reduction in superoxide production. Different letters indicate significant differences (*P* < 0.001; Tukey’s test). n = 5.

**Figure S2. *In vivo* characterisation of roGFP2-Orp1 targeted to various subcellular compartments in Arabidopsis pollen tubes.** (A) Subcellular localisation of roGFP2-Orp1 targeted to the cytosol, mitochondria, plastids, peroxisomes and nuclei. Representative images show fluorescence emission detected at 505–535 nm after excitation at 488 nm. Scale bar = 10 μm. (B) Initial oxidation states of roGFP2-Orp1 in the cytosol, mitochondria, plastids, peroxisomes and nuclei. Different letters indicate significant differences (*P* < 0.001) based on Tukey’s test. n ≥ 17. (C-G) Dynamic range characterisation of roGFP2-Orp1 oxidation and reduction in response to H_2_O_2_ and DTT in various subcellular compartments. Pollen tubes expressing roGFP2-Orp1 targeted to the cytosol (C), mitochondria (D), plastids (E), nuclei (F) and peroxisomes (G) were treated with 15 mM H_2_O_2_ or 10 mM DTT. The 405/488 nm excitation fluorescence ratio (R_405/488_) was measured *via* live-cell imaging and normalised to the initial mean R_405/488_. The dynamic range (δ) of roGFP2-Orp1 in each compartment was calculated using the R405/488 after oxidation (H_2_O_2_ treatment) and reduction (DTT treatment). n ≥ 10.

**Figure S3. Time-lapse imaging of mitochondria- and plastid-targeted roGFP2-Orp1 in Arabidopsis pollen tubes before and after SI induction.** Representative time-lapse images showing roGFP2-Orp1 fluorescence in mitochondria (A) or plastids (B) of Arabidopsis pollen tubes, excited at 488 nm and detected at 505-535 nm (upper panel) before and after SI induction. The lower panel shows F-actin labelled with lifeact-mRuby2. The colour gradient indicates the fluorescence intensity, which decreases after SI induction due to oxidation of the sensor. Scale bar = 10 μm.

**Figure S4. SI does not impair the reduction capacity of roGFP2-Orp1 in the cytosol and mitochondria of Arabidopsis pollen tubes.** Quantification of 405/488 nm fluorescence ratio (R_405/488_) of roGFP2-Orp1 targeted to the cytosol (A), mitochondria (B) and plastids (C) in untreated (UT) pollen tubes or after 5 minutes of H_2_O_2_ treatment, with and without a preceding 1.5-minute SI induction. Sample sizes: A, n ≥ 10. B, n = 15. C, n ≥ 10. Different letters indicate significant differences based on Tukey’s test (*P* < 0.001).

**Figure S5. Antimycin A induces rapid decrease in mitochondrial membrane potential in Arabidopsis pollen tubes.** Quantification of TMRM fluorescence in mitochondria after treatment with antimycin A (AA). Different letters indicate significant (*P* < 0.05) differences based on Tukey’s test (compact letter display). n = 24, 10, 15, 16, 15, 10, 18.

**Figure S6. Manipulation of [Ca^2+^]_cyt_ in Arabidopsis pollen tubes by A23187 and GdCl_3_.** Quantification of YC3.6 fluorescence ratio (R_Venus/CFP_) in pollen tubes treated with growth medium (GM), PrsS_1_ (SI), calcium ionophore (A23187), or PrsS_1_ in combination with GdCl_3_ (SI + GdCl_3_). n ≥ 9. R_Venus/CFP_ values were normalised to the mean of GM-treated samples within the first 10 minutes of treatment. Different letters indicate significant differences based on Tukey’s test (*P* < 0.001).

**Figure S7. GAPDH activities in *Papaver* pollen tubes after SI induction and H_2_O_2_ treatment.** GAPDH activities were measured in *Papaver* pollen tubes 15 minutes after treatment with 2.5 mM H₂O₂ (\**P* < 0.05, Mann-Whitney U test, n = 3) and 30 minutes after SI induction (\**P* < 0.05, Mann-Whitney U test, n = 4). All GAPDH activities were normalised to the mean values of untreated (UT) or control samples.

**Figure S8. Changes in TCA cycle metabolite levels in Arabidopsis pollen tubes following SI induction.** Metabolite levels of malate (A), succinate (B) and citrate/isocitrate (C) decreased over the 0-120-minute period following SI induction, though not significantly. In contrast, 2-oxoglutarate (D) showed a significant decrease starting 15 minutes after SI induction. Different letters indicate significant differences based on Tukey’s test (*P* < 0.001, n = 4).

**Table S1. Transgenic Arabidopsis lines used in this study.**

**Table S2. Oligonucleotides, targeting peptide sequences and Golden Gate modules used for transgene construction.** A. List of oligonucleotides used in this study. F, forward primers. R, reverse primers. Bold letters indicate target-binding sequences. B. Targeting peptide sequences fused to roGFP2-Orp1 for subcellular localisation; C. Golden Gate Assembly modules used in this study.

**Table S3. Multiple reaction monitoring (MRM) transitions for LC-triple quadrupole MS/MS analysis of TCA cycle acids and glutathione.**

## Notes

### Competing Interest Statement

The authors have declared no competing interest.

## References

Arnaud D, Deeks MJ, Smirnoff N. Organelle-targeted biosensors reveal distinct oxidative events during pattern-triggered immune responses. Plant Physiology. 2023:191(4):2551–2569. 10.1093/plphys/kiac603

Bénit P, Chrétien D, Porceddu M, Yanicostas C, Rak M, Rustin P. An Effective, Versatile, and Inexpensive Device for Oxygen Uptake Measurement. J Clin Med. 2017:6(6):58. 10.3390/jcm6060058

Bienert GP, Chaumont F. Aquaporin-facilitated transmembrane diffusion of hydrogen peroxide. Biochim Biophys Acta. 2014:1840(5):1596–1604. 10.1016/j.bbagen.2013.09.017

Boisson-Dernier A, Lituiev DS, Nestorova A, Franck CM, Thirugnanarajah S, Grossniklaus U. ANXUR receptor-like kinases coordinate cell wall integrity with growth at the pollen tube tip *via* NADPH oxidases. Plos Biology. 2013:11(11):e1001719. 10.1371/journal.pbio.1001719

Bosch M, Franklin-Tong, VE. Temporal and spatial activation of caspase-like enzymes induced by self-incompatibility in *Papaver* pollen. Proc Natl Acad Sci U S A. 2007:104(46):18327–18332. 10.1073/pnas.0705826104

Bosch M, Franklin-Tong VE. Regulating PCD in plant cells: Intracellular acidification plays a pivotal role together with calcium signaling. Plant Cell. 2024:36(11):4692–4702. 10.1093/plcell/koae245

Boussardon C, Simon M, Carrie C, Fuszard M, Meyer EH, Budar F, Keech O. The atypical proteome of mitochondria from mature pollen grains. Curr Biol. 2025:35(4):776–787. 10.1016/j.cub.2024.12.037

Carrie C, Venne AS, Zahedi RP, Soll J. Identification of cleavage sites and substrate proteins for two mitochondrial intermediate peptidases in *Arabidopsis thaliana*. J Exp Bot. 2015:66(9):2691–2708. 10.1093/jxb/erv064

Chaumont F, de Castro Silva Filho M, Thomas D, Leterme S, Boutry M. Truncated presequences of mitochondrial F1-ATPase beta subunit from *Nicotiana plumbaginifolia* transport CAT and GUS proteins into mitochondria of transgenic tobacco. Plant Mol Biol. 1994:24(4):631–641. 10.1007/bf00023559

Clough SJ, Bent AF. Floral dip: a simplified method for Agrobacterium-mediated transformation of *Arabidopsis thaliana*. The Plant Journal. 1998:16(6):735–743. 10.1046/j.1365-313x.1998.00343.x

Curtis MJ, Wolpert TJ. The victorin-induced mitochondrial permeability transition precedes cell shrinkage and biochemical markers of cell death, and shrinkage occurs without loss of membrane integrity. The Plant Journal. 2004:38(2):244–259. 10.1111/j.1365-313X.2004.02040.x

de Graaf BH, Vatovec S, Juarez-Diaz JA, Chai L, Kooblall K, Wilkins KA., Zou H, Forbes T, Franklin FC, Franklin-Tong VE. The Papaver self-incompatibility pollen *S*-determinant, PrpS, functions in *Arabidopsis thaliana*. Curr Biol. 2012:22(2):154–159. 10.1016/j.cub.2011.12.006

Duchen MR. Mitochondria and calcium: from cell signalling to cell death. J Physiol. 2000:529:57–68. 10.1111/j.1469-7793.2000.00057.x

Dumont S, Rivoal J. Consequences of Oxidative Stress on Plant Glycolytic and Respiratory Metabolism. Frontiers in Plant Science. 2019:10:166. 10.3389/fpls.2019.00166

Emes MJ, Neuhaus HE. Metabolism and transport in non-photosynthetic plastids. J Exp Bot. 1997:48(12):1995–2005. 10.1093/jxb/48.12.1995

Engler C, Youles M, Gruetzner R, Ehnert TM, Werner S, Jones JD, Patron NJ, Marillonnet S. A golden gate modular cloning toolbox for plants. ACS Synth Biol. 2014:3(11):839–843. 10.1021/sb4001504

Exposito-Rodriguez M, Laissue PP, Yvon-Durocher G, Smirnoff N, Mullineaux PM. (2017) Photosynthesis-dependent H_2_O_2_ transfer from chloroplasts to nuclei provides a high-light signalling mechanism. Nature Communications 8: 4. 10.1038/s41467-017-00074-w

Foote HC, Ride JP, Franklin-Tong, VE, Walker EA, Lawrence MJ, Franklin FC. Cloning and expression of a distinctive class of self-incompatibility (*S*) gene from *Papaver rhoeas* L. Proc Natl Acad Sci U S A. 1994:91(6):2265–2269. 10.1073/pnas.91.6.22

Foreman J, Demidchik V, Bothwell JHF, Mylona P, Miedema H, Torres MA, Linstead P, Costa S, Brownlee C, Jones JDG, et al. Reactive oxygen species produced by NADPH oxidase regulate plant cell growth. Nature. 2003:422(6930):442-446. 10.1038/nature01485

Franklin-Tong VE, Ride JP, Read ND, Trewavas AJ, Franklin FC. The self-incompatibility response in *Papaver rhoeas* is mediated by cytosolic free calcium. The Plant Journal. 1993:4(1):163–177. 10.1046/j.1365-313X.1993.04010163.x

Franklin-Tong, VE, Bosch M. Plant biology: Stigmatic ROS decide whether pollen is accepted or rejected. Current Biology. 2021:31(14):R904–R906. 10.1016/j.cub.2021.05.034

Geitmann A, Snowman BN, Emons AM, Franklin-Tong VE. Alterations in the actin cytoskeleton of pollen tubes are induced by the self-incompatibility reaction in *Papaver rhoeas*. Plant Cell. 2000:12(7):1239–1251. 10.1105/tpc.12.7.1239

Geitmann A, Franklin-Tong VE, Emons AC. The self-incompatibility response in *Papaver rhoeas* pollen causes early and striking alterations to organelles. Cell Death Differ. 2004:11(8):812–822. 10.1038/sj.cdd.4401424

Gibon Y, Blaesing OE, Hannemann J, Carillo P, Höhne M, Hendriks JH, Palacios P, Cross J, Selbig J, Stitt M. A Robot-based platform to measure multiple enzyme activities in Arabidopsis using a set of cycling assays: comparison of changes of enzyme activities and transcript levels during diurnal cycles and in prolonged darkness. Plant Cell. 2004:16(12):3304–3325. 10.1105/tpc.104.025973

Gutscher M, Sobotta MC, Wabnitz GH, Ballikaya S, Meyer AJ, Samstag Y, Dick TP. Proximity-based protein thiol oxidation by H_2_O_2_-scavenging peroxidases. Journal of Biological Chemistry. 2009:284(46):31532–31540. 10.1074/jbc.M109.059246

Haque T, Eaves DJ, Lin Z, Zampronio CG, Cooper HJ, Bosch M, Smirnoff N, Franklin-Tong, VE. Self-Incompatibility Triggers Irreversible Oxidative Modification of Proteins in Incompatible Pollen. Plant Physiology. 2020:183(3):1391–1404. 10.1104/pp.20.00066

Hempel SL, Buettner GR, O’Malley YQ, Wessels DA, Flaherty DM. Dihydrofluorescein diacetate is superior for detecting intracellular oxidants: comparison with 2’,7’-dichlorodihydrofluorescein diacetate, 5(and 6)-carboxy-2’,7’-dichlorodihydrofluorescein diacetate, and dihydrorhodamine 123. Free Radic Biol Med. 1999:27(1-2):146–159. 10.1016/s0891-5849(99)00061-1

Hepler PK, Lovy-Wheeler A, McKenna ST, Kunkel JG. Ions and Pollen Tube Growth. In R. Malhó (eds), The Pollen Tube: A Cellular and Molecular Perspective (pp. 47–69). Plant Cell Monographs, vol 3. Springer, Berlin, Heidelberg. 10.1007/7089_043

Hillion M, Imber M, Pedre B, Bernhardt J, Saleh M, Loi VV., Maaß S, Becher D, Rosado LA, Adrian L, et al. The glyceraldehyde-3-phosphate dehydrogenase GapDH of Corynebacterium diphtheriae is redox-controlled by protein S-mycothiolation under oxidative stress. Scientific Reports. 2017:7(1):5020. 10.1038/s41598-017-05206-2

Holdaway-Clarke TL, Hepler PK. Control of pollen tube growth: role of ion gradients and fluxes. New Phytologist. 2003:159(3):539–563. 10.1046/j.1469-8137.2003.00847.x

Huang SB, Van Aken O, Schwarzlander M, Belt K, Millar AH. The Roles of Mitochondrial Reactive Oxygen Species in Cellular Signaling and Stress Response in Plants. Plant Physiology. 2016:171(3):1551–1559. 10.1104/pp.16.00166

Kacprzyk J, Burke R, Armengot L, Coppola M, Tattrie SB, Vahldick H, Bassham DC, Bosch M, Brereton NJB, Cacas J-L, et al. Roadmap for the next decade of plant programmed cell death research. New Phytol. 2024:242(5):1865–1875. 10.1111/nph.19709

Kaya H, Nakajima R, Iwano M, Kanaoka MM, Kimura S, Takeda S, Kawarazaki T, Senzaki E, Hamamura Y, Higashiyama T, et al. Ca^2+^-activated reactive oxygen species production by Arabidopsis RbohH and RbohJ is essential for proper pollen tube tip growth. Plant Cell. 2014:26(3):1069–1080. 10.1105/tpc.113.120642

Kaya H, Iwano M, Takeda S, Kanaoka M M, Kimura S, Abe M, Kuchitsu K. Apoplastic ROS production upon pollination by RbohH and RbohJ in Arabidopsis. Plant Signal Behav. 2015:10(2):e989050. 10.4161/15592324.2014.989050

Khan K, Tran HC, Mansuroglu B, Onsell P, Buratti S, Schwarzlander M, Costa A, Rasmusson AG, Van Aken O. Mitochondria-derived reactive oxygen species are the likely primary trigger of mitochondrial retrograde signaling in *Arabidopsis*. Curr Biol. 2024:34(2):327–342. 10.1016/j.cub.2023.12.005

Lampropoulos A, Sutikovic Z, Wenzl C, Maegele I, Lohmann JU, Forner J. GreenGate - A Novel, Versatile, and Efficient Cloning System for Plant Transgenesis. PLOS ONE. 2013:8(12):e83043. 10.1371/journal.pone.0083043

Lassig R, Gutermuth T, Bey TD, Konrad KR, Romeis T. Pollen tube NAD(P)H oxidases act as a speed control to dampen growth rate oscillations during polarized cell growth. Plant J. 2014:78(1):94–106. 10.1111/tpj.12452

Lin Z, Eaves DJ, Sanchez-Moran E, Franklin FC, Franklin-Tong VE. The *Papaver rhoeas S* determinants confer self-incompatibility to Arabidopsis thaliana in planta. Science. 2015:350(6261):684–687. 10.1126/science.aad2983

Lin Z, Xie F, Trivino M, Karimi M, Bosch M, Franklin-Tong VE, Nowack MK. Ectopic Expression of a Self-Incompatibility Module Triggers Growth Arrest and Cell Death in Vegetative Cells. Plant Physiol. 2020:183(4):1765–1779. 10.1104/pp.20.00292

Liu J, Lim SL, Zhong JY, Lim BL. Bioenergetics of pollen tube growth in *Arabidopsis thaliana* revealed by ratiometric genetically encoded biosensors. Nat Commun. 2022:13(1):7822. 10.1038/s41467-022-35486-w

Loro G, Drago I, Pozzan T, Schiavo FL, Zottini M, Costa A. Targeting of Cameleons to various subcellular compartments reveals a strict cytoplasmic/mitochondrial Ca^2+^ handling relationship in plant cells. The Plant Journal. 2012:71:1–13. 10.1111/j.1365-313X.2012.04968.x

Maxwell DP, Wang Y, McIntosh L. The alternative oxidase lowers mitochondrial reactive oxygen production in plant cells. Proc Natl Acad Sci U S A. 1999:96(14):8271–8276. 10.1073/pnas.96.14.8271

Mittler R, Zilinskas BA. Purification and characterization of pea cytosolic ascorbate peroxidase. Plant Physiol. 1991: 97: 962–968. 10.1104/pp.97.3.962.

Mittler R, Zandalinas SI, Fichman Y, Van Breusegem F. Reactive oxygen species signalling in plant stress responses. Nature Reviews Molecular Cell Biology. 2022:23(10):663–679. 10.1038/s41580-022-00499-2

Moller, IM. Plant mitochondria and oxidative stress: Electron transport, NADPH turnover, and metabolism of reactive oxygen species. Annual Review of Plant Physiology and Plant Molecular Biology. 2001:52:561–591. 10.1146/annurev.arplant.52.1.561

Moller IM, Rasmusson AG, Van Aken O. Plant mitochondria - past, present and future. Plant Journal. 2021:108(4):912–959. 10.1111/tpj.15495

Mur LAJ, Kenton P, Lloyd AJ, Ougham H, Prats E. The hypersensitive response; the centenary is upon us but how much do we know? Journal of Experimental Botany. 2008:59(3):501–520. 10.1093/jxb/erm239

Nietzel T, Elsässer M, Ruberti C, Steinbeck J, Ugalde JM, Fuchs P, Wagner S, Ostermann L, Moseler A, Lemke P, et al. The fluorescent protein sensor roGFP2-Orp1 monitors *in vivo* H_2_O_2_ and thiol redox integration and elucidates intracellular H_2_O_2_ dynamics during elicitor-induced oxidative burst in Arabidopsis. New Phytologist. 2019:221(3):1649–1664. 10.1111/nph.15550

Noctor G, Foyer CH. Intracellular Redox Compartmentation and ROS-Related Communication in Regulation and Signaling. Plant Physiology. 2016:171(3):1581–1592. 10.1104/pp.16.00346

Obermeyer G, Fragner L, Lang V, Weckwerth W. Dynamic Adaption of Metabolic Pathways during Germination and Growth of Lily Pollen Tubes after Inhibition of the Electron Transport Chain. Plant Physiology. 2013:162(4):1822–1833. 10.1104/pp.113.219857

Ocampo-Hernández B, Gutiérrez Mireles ER, Gutiérrez-Aguilar M. Morphology and permeability transitions in plant mitochondria: Different aspects of the same event? Biochim Biophys Acta Bioenerg. 2022:1863(7):148586. 10.1016/j.bbabio.2022.148586

Page MT, McCormac AC, Smith AG, Terry MJ. Singlet oxygen initiates a plastid signal controlling photosynthetic gene expression. New Phytologist. 2017: 213: 1168–1180. 10.1111/nph.14223

Perry SW, Norman JP, Barbieri J, Brown EB, Gelbard HA. Mitochondrial membrane potential probes and the proton gradient: a practical usage guide. BioTechniques. 2011:50(2):98–115. 10.2144/000113610

Pierson ES, Miller DD, Callaham DA, van Aken J, Hackett G, Hepler PK. Tip-Localized Calcium Entry Fluctuates during Pollen Tube Growth. Developmental Biology. 1996:174(1):160–173. 10.1006/dbio.1996.0060

Postiglione AE, Muday GK. Abscisic acid increases hydrogen peroxide in mitochondria to facilitate stomatal closure. Plant Physiology 2023: 192: 469–487. 10.1093/plphys/kiac601

Potocký M, Eliáš M, Profotová B, Novotná Z, Valentová O, Žárský V. Phosphatidic acid produced by phospholipase D is required for tobacco pollen tube growth. Planta. 2003:217(1):122–130. 10.1007/s00425-002-0965-4

Potocký M, Jones MA, Bezvoda R, Smirnoff N, Žárský V. Reactive oxygen species produced by NADPH oxidase are involved in pollen tube growth. New Phytologist. 2007:174(4):742–751. 10.1111/j.1469-8137.2007.02042.x

Potocký M, Pejchar P, Gutkowska M, Jiménez-Quesada MJ, Potocká A, de Dios Alché J, Kost B, Žárský V. NADPH oxidase activity in pollen tubes is affected by calcium ions, signaling phospholipids and Rac/Rop GTPases. Journal of Plant Physiology. 2012:169(16):1654–1663. 10.1016/j.jplph.2012.05.014

Rattanawong K, Koiso N, Toda E, Kinoshita A, Tanaka M, Tsuji H, Okamoto T. Regulatory functions of ROS dynamics *via* glutathione metabolism and glutathione peroxidase activity in developing rice zygote. The Plant Journal. 2021:108(4):1097–1115. 10.1111/tpj.15497

Renato M, Boronat A, Azcon-Bieto J. Respiratory processes in non-photosynthetic plastids. Front Plant Sci. 2015:6:496. 10.3389/fpls.2015.00496

Rodrigues O, Reshetnyak G, Grondin A, Saijo Y, Leonhardt N, Maurel C, Verdoucq L. Aquaporins facilitate hydrogen peroxide entry into guard cells to mediate ABA- and pathogen-triggered stomatal closure. Proceedings of the National Academy of Sciences. 2017:114(34):9200–9205. 10.1073/pnas.1704754114

Rounds CM, Lubeck E, Hepler PK, Winship LJ. Propidium iodide competes with Ca(2+) to label pectin in pollen tubes and Arabidopsis root hairs. Plant Physiology. 2011:157(1):175–187. 10.1104/pp.111.182196

Scaduto RC, Grotyohann LW. Measurement of Mitochondrial Membrane Potential Using Fluorescent Rhodamine Derivatives. Biophysical Journal. 1999:76(1):469–477. 10.1016/S0006-3495(99)77214-0

Schindelin J, Arganda-Carreras I, Frise E, Kaynig V, Longair M, Pietzsch T, Preibisch S, Rueden C, Saalfeld S, Schmid B, et al. Fiji: an open-source platform for biological-image analysis. Nature Methods. 2012:9(7):676–682. 10.1038/nmeth.2019

Schneider M, Knuesting J, Birkholz O, Heinisch JJ, Scheibe R. Cytosolic GAPDH as a redox-dependent regulator of energy metabolism. BMC Plant Biol. 2018:18(1):184. 10.1186/s12870-018-1390-6

Scott I, Logan DC. Mitochondrial morphology transition is an early indicator of subsequent cell death in Arabidopsis. New Phytologist. 2008:177(1):90–101. 10.1111/j.1469-8137.2007.02255.x

Smirnoff N, Arnaud D. Hydrogen peroxide metabolism and functions in plants. New Phytologist. 2019:221(3):1197–1214. 10.1111/nph.15488

Snowman BN, Kovar DR, Shevchenko G, Franklin-Tong VE, Staiger CJ. Signal-mediated depolymerization of actin in pollen during the self-incompatibility response. Plant Cell. 2022:14(10):2613–2626.

Thomas SG, Franklin-Tong VE. Self-incompatibility triggers programmed cell death in *Papaver* pollen. Nature. 2004:429(6989):305–309. 10.1038/nature02540

Torres MA, Dangl JL, Jones JDG. Arabidopsis gp91(phox) homologues *AtrbohD* and *AtrbohF* are required for accumulation of reactive oxygen intermediates in the plant defense response. Proceedings of the National Academy of Sciences of the United States of America 2002: 99: 517–522. 10.1073pnas.012452499

Ugalde JM, Fuchs P, Nietzel T, Cutolo EA, Homagk M, Vothknecht UC, Holuigue L, Schwarzländer M, Müller-Schüssele SJ, Meyer AJ. Chloroplast-derived photo-oxidative stress causes changes in H_2_O_2_ and E_GSH_ in other subcellular compartments. Plant Physiology. 2021:186(1):125–141. 10.1093/plphys/kiaa095

Van Aken O, Van Breusegem F. Licensed to Kill: Mitochondria, Chloroplasts, and Cell Death. Trends Plant Sci. 2015:20(11):754–766. 10.1016/j.tplants.2015.08.002

Wagner S, Steinbeck J, Fuchs P, Lichtenauer S, Elsässer M, Schippers JHM, Nietzel T, Ruberti C, Van Aken O, Meyer AJ, et al. Multiparametric real-time sensing of cytosolic physiology links hypoxia responses to mitochondrial electron transport. New Phytologist. 2019:224(4):1668–1684. 10.1111/nph.16093

Walvekar A, Rashida Z, Maddali H, Laxman S. A versatile LC-MS/MS approach for comprehensive, quantitative analysis of central metabolic pathways. Wellcome Open Res. 2018:3:122. 10.12688/wellcomeopenres.14832.1

Wang L, Lin Z, Triviño M, Nowack MK, Franklin-Tong VE, Bosch M. Self-incompatibility in *Papaver* pollen: programmed cell death in an acidic environment. J Exp Bot. 2019:70(7):2113–2123. 10.1093/jxb/ery406

Wang L, Triviño M, Lin Z, Carli J, Eaves DJ, Van Damme D, Nowack MK, Franklin-Tong VE, Bosch M. New opportunities and insights into *Papaver* self-incompatibility by imaging engineered Arabidopsis pollen. J Exp Bot. 2020:71(8):2451–2463. 10.1093/jxb/eraa092

Wang L, Lin Z, Carli J, Gladala-Kostarz A, Davies JM, Franklin-Tong VE, Bosch M. ATP depletion plays a pivotal role in self-incompatibility, revealing a link between cellular energy status, cytosolic acidification and actin remodelling in pollen tubes. New Phytol. 2022:236(5):1691–1707. 10.1111/nph.18350

Wang J, Bollier N, Buono RA, Vahldick H, Lin Z, Feng Q, Hudecek R, Jiang Q, Mylle E, Van Damme D, Nowack MK. A developmentally controlled cellular decompartmentalization process executes programmed cell death in the Arabidopsis root cap. The Plant Cell. 2024:36(4):941–962. 10.1093/plcell/koad308

Weber E, Gruetzner R, Werner S, Engler C, Marillonnet S. Assembly of Designer TAL Effectors by Golden Gate Cloning. PLOS ONE. 2011:6(5):e19722. 10.1371/journal.pone.0019722

Wendt S, Johnson S, Weilinger NL, Groten C, Sorrentino S, Frew J, Yang L, Choi HB, Nygaard HB, MacVicar BA. Simultaneous imaging of redox states in dystrophic neurites and microglia at Aβ plaques indicate lysosome accumulation not microglia correlate with increased oxidative stress. Redox Biol. 2022:56:102448. 10.1016/j.redox.2022.102448

Weterings K, Reijnen W, Wijn G, van de Heuvel K, Appeldoorn N, de Kort G, van Herpen M, Schrauwen J, Wullems G. Molecular characterization of the pollen-specific genomic clone NTPg303 and in situ localization of expression. Sexual Plant Reproduction. 1995:8(1):11–17. 10.1007/BF00228757

Wheeler MJ, de Graaf BH, Hadjiosif N, Perry RM, Poulter NS, Osman K, Vatovec S, Harper A, Franklin FC, Franklin-Tong VE. Identification of the pollen self-incompatibility determinant in *Papaver rhoeas*. Nature. 2009:459(7249):992–995. 10.1038/nature08027

Wilkins KA, Bancroft J, Bosch M, Ings J, Smirnoff N, Franklin-Tong VE. Reactive oxygen species and nitric oxide mediate actin reorganization and programmed cell death in the self-incompatibility response of *Papaver*. Plant Physiology. 2011:156(1):404–416. 10.1104/pp.110.167510

Wilkins KA, Bosch M, Haque T, Teng N, Poulter NS, Franklin-Tong VE. Self-incompatibility-induced programmed cell death in field poppy pollen involves dramatic acidification of the incompatible pollen tube cytosol. Plant Physiology. 2015:167(3):766–779. 10.1104/pp.114.252742

Xie HT, Wan ZY, Li S, Zhang Y. Spatiotemporal Production of Reactive Oxygen Species by NADPH Oxidase Is Critical for Tapetal Programmed Cell Death and Pollen Development in *Arabidopsis*. Plant Cell. 2014:26(5):2007–2023. 10.1105/tpc.114.125427

Zhang L, Huang J, Su S, Wei X, Yang L, Zhao H, Yu J, Wang J, Hui J, Hao S, et al. FERONIA receptor kinase-regulated reactive oxygen species mediate self-incompatibility in *Brassica rapa*. Current Biology. 2021:31(14):3004–3016.e3004. 10.1016/j.cub.2021.04.060

