## Supplemental Figures and Tables for "*Papaver S*-determinants trigger an integrated network of mitochondrially derived ROS and disruption of energy metabolism in incompatible pollen tubes"

**Figure S1. Disruption of tip-localized NADPH oxidase activity in *Papaver* pollen tubes following SI induction.**

**Figure S2. *In vivo* characterisation of roGFP2-Orp1 targeted to various subcellular compartments in Arabidopsis pollen tubes.**

**Figure S3. Time-lapse imaging of mitochondria- and plastid-targeted roGFP2-Orp1 in Arabidopsis pollen tubes before and after SI induction.**

**Figure S4. SI does not impair the reduction capacity of roGFP2-Orp1 in the cytosol and mitochondria of Arabidopsis pollen tubes.**

**Figure S5. Antimycin A induces rapid decrease in mitochondrial membrane potential in Arabidopsis pollen tubes.**

**Figure S6. Manipulation of $[Ca^{2+}]_{cyt}$ in Arabidopsis pollen tubes by A23187 and $GdCl_3$.**

**Figure S7. GAPDH activities in *Papaver* pollen tubes after SI induction and $H_2O_2$ treatment.**

**Figure S8. Changes in TCA cycle metabolite levels in Arabidopsis pollen tubes following SI induction.**

**Table S1. Transgenic Arabidopsis lines used in this study.**

**Table S2. Oligonucleotides, targeting peptide sequences and Golden Gate modules used for transgene construction.**

**Table S3. Multiple reaction monitoring (MRM) transitions for LC-triple quadrupole MS/MS analysis of TCA cycle acids and glutathione.**

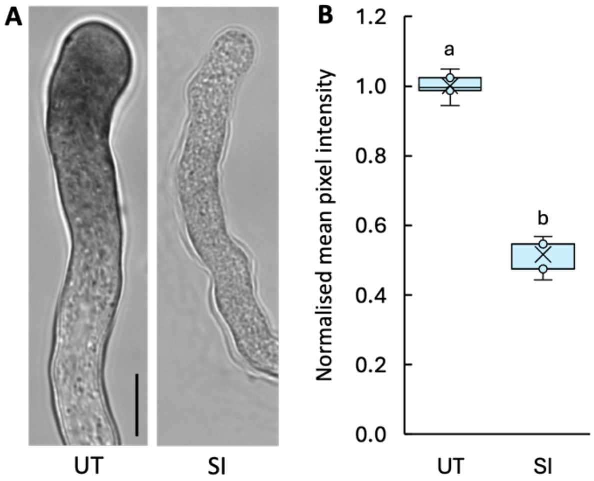


**Figure S1. Disruption of tip-localized NADPH oxidase activity in *Papaver* pollen tubes following SI induction.** (A) Nitroblue tetrazolium (NBT) staining was used to detect superoxide production at the tips of *Papaver* pollen tubes during the SI response. Representative images show an untreated (UT) pollen tube and a pollen tube treated with $PsS_1$ (SI) for 15 min. NBT staining was significantly reduced after SI induction. Scale bar = 10 µm. (B) Quantification of NBT staining intensity at pollen tube tips. Reduced NBT staining reflects decreased pixel intensity, indicating a reduction in superoxide production. Different letters indicate significant differences ($P < 0.001$; Tukey's test). n = 5.

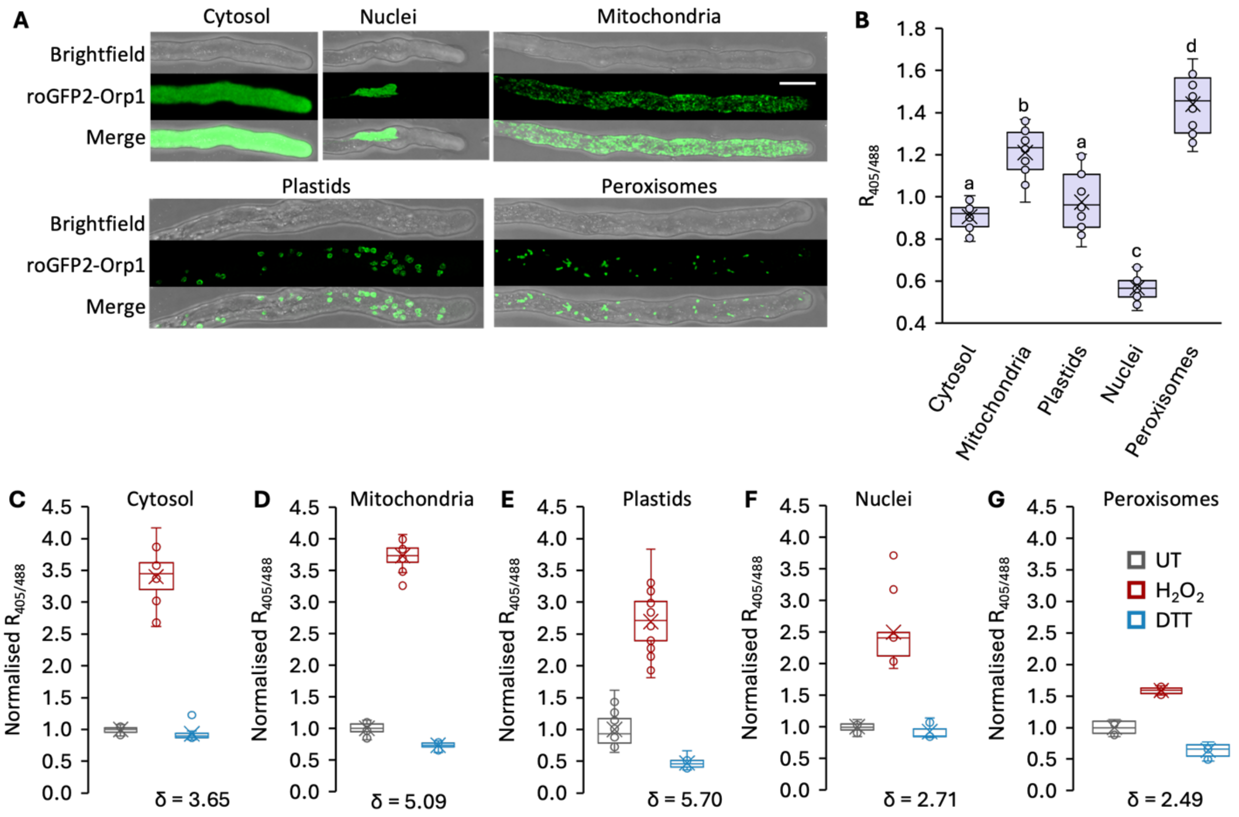


**Figure S2. *In vivo* characterisation of roGFP2-Orp1 targeted to various subcellular compartments in Arabidopsis pollen tubes.** (A) Subcellular localisation of roGFP2-Orp1 targeted to the cytosol, mitochondria, plastids, peroxisomes and nuclei. Representative images show fluorescence emission detected at 505–535 nm after excitation at 488 nm. Scale bar = 10 µm. (B) Initial oxidation states of roGFP2-Orp1 in the cytosol, mitochondria, plastids, peroxisomes and nuclei. Different letters indicate significant differences ($P < 0.001$) based on Tukey's test $n \geq 17$. (C-G) Dynamic range characterisation of roGFP2-Orp1 oxidation and reduction in response to $H_2O_2$ and DTT in various subcellular compartments. Pollen tubes expressing roGFP2-Orp1 targeted to the cytosol (C), mitochondria (D), plastids (E), nuclei (F) and peroxisomes (G) were treated with 15 mM $H_2O_2$ or 10 mM DTT. The 405/488 nm excitation fluorescence ratio ($R_{405/488}$) was measured via live-cell imaging and normalised to the initial mean $R_{405/488}$. The dynamic range (δ) of roGFP2-Orp1 in each compartment was calculated using the $R_{405/488}$ after oxidation ($H_2O_2$ treatment) and reduction (DTT treatment). $n \geq 10$.

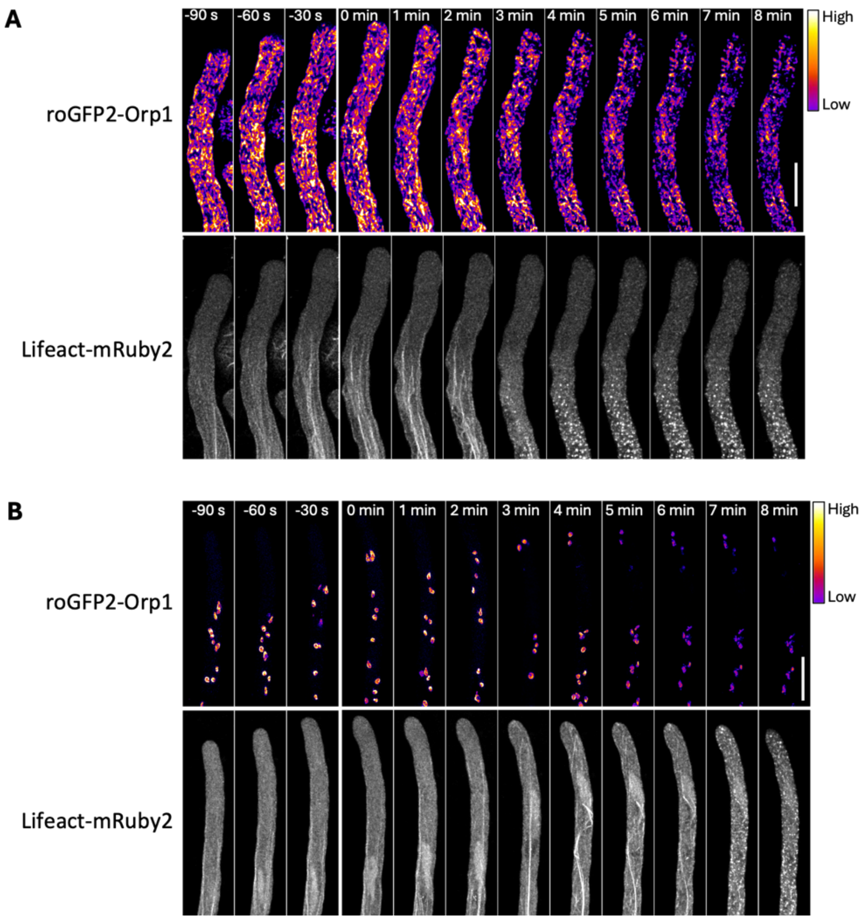


**Figure S3. Time-lapse imaging of mitochondria- and plastid-targeted roGFP2-Orp1 in Arabidopsis pollen tubes before and after SI induction.** Representative time-lapse images showing roGFP2-Orp1 fluorescence in mitochondria (A) or Plastids (B) of Arabidopsis pollen tubes, excited at 488 nm and detected at 505-535 nm (upper panel) before and after SI induction. The lower panel shows F-actin labelled with lifeact-mRuby2. The colour gradient indicates the fluorescence intensity, which decreases after SI induction due to oxidation of the sensor. Scale bar = 10 µm.

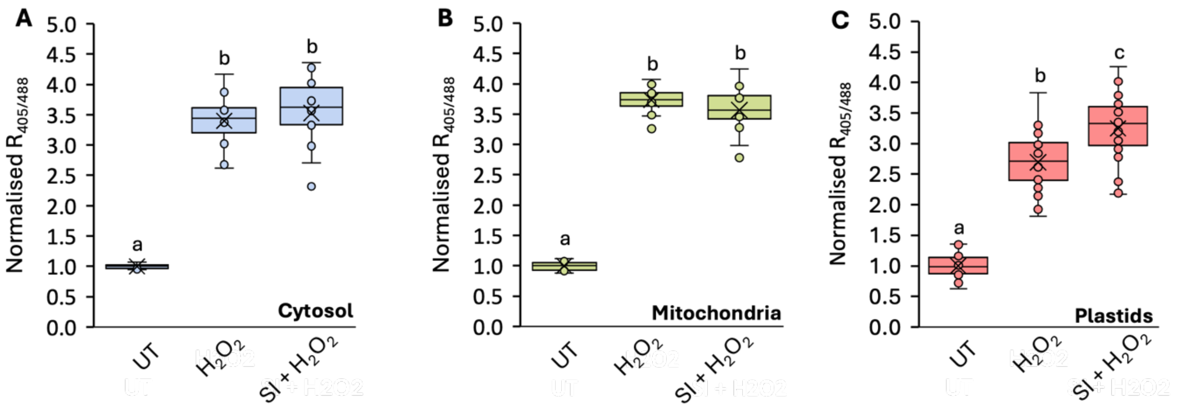


**Figure S4. SI does not impair the reduction capacity of roGFP2-Orp1 in the cytosol and mitochondria of Arabidopsis pollen tubes.** Quantification of 405/488 nm fluorescence ratio ($R_{405/488}$) of roGFP2-Orp1 targeted to the cytosol (A), mitochondria (B) and plastids (C) in untreated (UT) pollen tubes or after 5 minutes of $H_2O_2$ treatment, with and without a preceding 1.5-minute SI induction. Sample sizes: A, n ≥ 10. B, n = 15. C, n ≥ 10. Different letters indicate significant differences based on Tukey's test ($P < 0.001$).

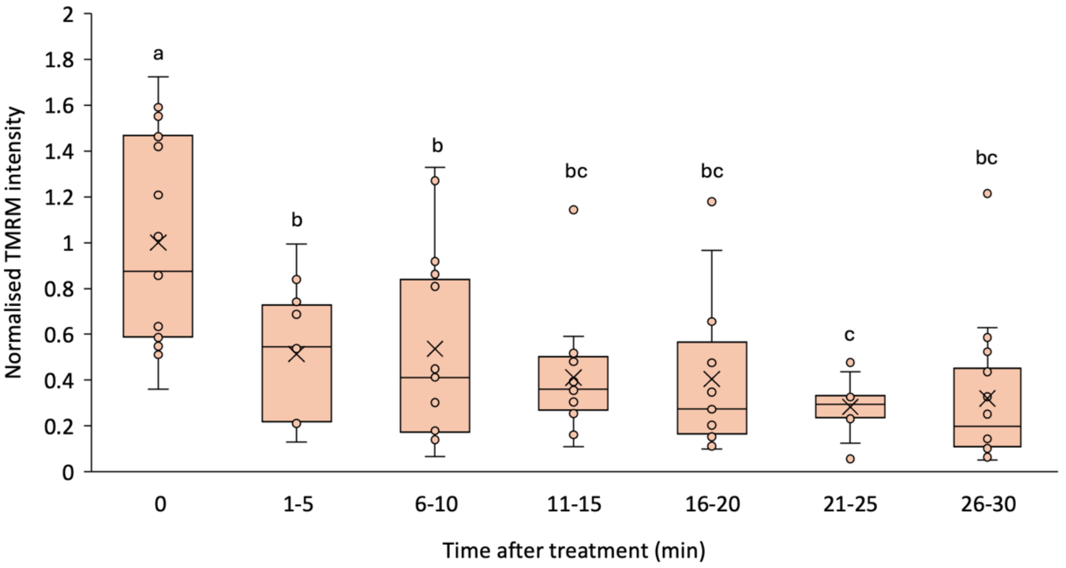


**Figure S5. Antimycin A induces rapid decrease in mitochondrial membrane potential in Arabidopsis pollen tubes.** Quantification of TMRM fluorescence in mitochondria after treatment with antimycin A (AA). Different letters indicate significant ($P < 0.05$) differences based on Tukey's test (compact letter display). n = 24, 10, 15, 16, 15, 10, 18.

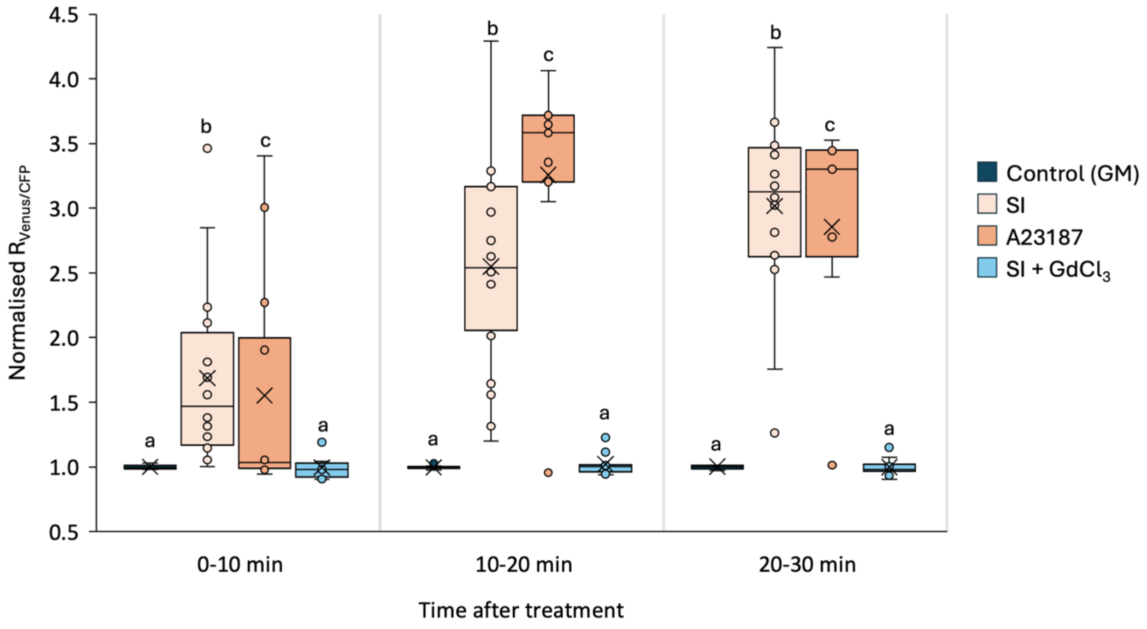


**Figure S6. Manipulation of $[Ca^{2+}]_{cyt}$ in Arabidopsis pollen tubes by A23187 and $GdCl_3$.** Quantification of YC3.6 fluorescence ratio ($R_{Venus/CFP}$) in pollen tubes treated with growth medium (GM), $PrsS_1$ (SI), calcium ionophore (A23187), or $PrsS_1$ in combination with $GdCl_3$ (SI + $GdCl_3$). n ≥ 9. $R_{Venus/CFP}$ values were normalised to the mean of GM-treated samples within the first 10 minutes of treatment. Different letters indicate significant differences based on Tukey's test ($P < 0.001$).

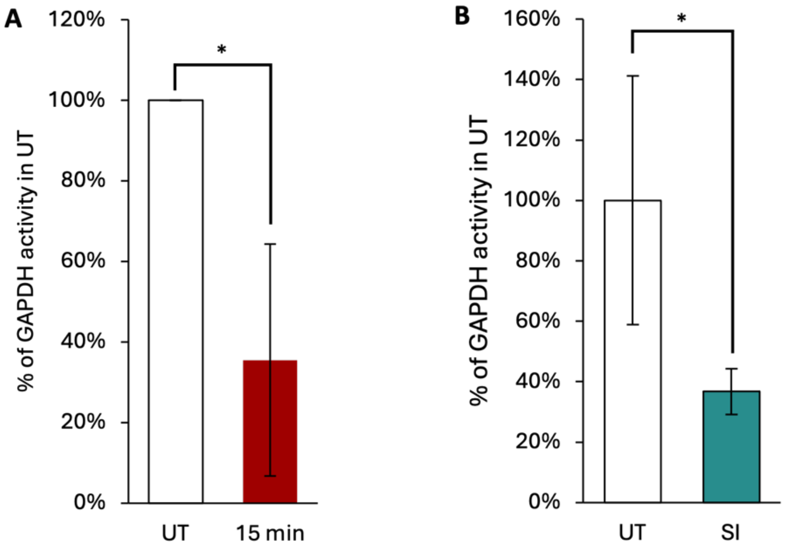


**Figure S7. GAPDH activities in *Papaver* pollen tubes after SI induction and $H_2O_2$ treatment.** GAPDH activities were measured in *Papaver* pollen tubes 15 minutes after treatment with 2.5 mM $H_2O_2$ (*$P < 0.05$, Mann-Whitney U test, n = 3) and 30 minutes after SI induction (*$P < 0.05$, Mann-Whitney U test, n = 4). All GAPDH activities were normalised to the mean values of untreated (UT) or control samples.

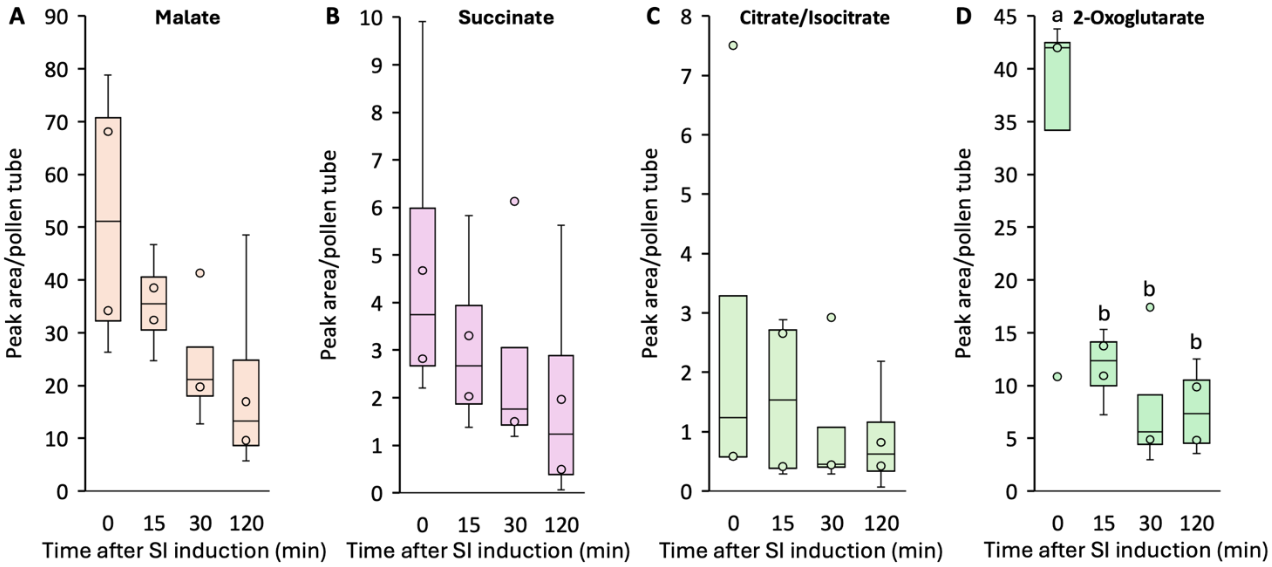


**Figure S8. Changes in TCA cycle metabolite levels in Arabidopsis pollen tubes following SI induction.** Metabolite levels of malate (A), succinate (B) and citrate/isocitrate (C) decreased over the 0-120 minute period following SI induction, though not significantly. In contrast, 2-oxoglutarate (D) showed a significant decrease starting 15 minutes after SI induction. Different letters indicate significant differences based on Tukey's test ($P < 0.001$, n = 4).

**Table S1. Transgenic Arabidopsis lines used in this study.**

| Lines | Constructs | Corresponding figures |
|---|---|---|
| 1 | *ProNTP303:roGFP2-Orp1, ProNTP303:Lifeact-mRuby2-ProNTP303:PrpS₁* | Figure 1, Figure S2, Figure 2A and 2F, Figure 3, Figure S4A, Figure 4A and 4C, Figure 6F and 6G, Figure 7A |
| 2 | *ProNTP303:Mito-roGFP2-Orp1, ProNTP303:Lifeact-mRuby2-ProNTP303:PrpS₁* | Figure S2, Figure 2B and 2G, Figure S3A, Figure S4B, Figure 4B, Figure 6D, Figure 7B |
| 3 | *ProNTP303:Chl-roGFP2-Orp1, ProNTP303:Lifeact-mRuby2-ProNTP303:PrpS₁* | Figure 2C and 2H, Figure S3B, Figure S4C, Figure 4D, Figure 6E, Figure 8C-E, Figure 9, Figure S8 |
| 4 | *ProNTP303:roGFP2-Orp1-SKL, ProNTP303:Lifeact-mRuby2-ProNTP303:PrpS₁* | Figure S2, Figure 2D |
| 5 | *ProNTP303:NLS-roGFP2-Orp1, ProNTP303:Lifeact-mRuby2-ProNTP303:PrpS₁* | Figure S2, Figure 2E |
| 6 | *ProNTP303:pHGFP-ProNTP303:PrpS₁ (Wang et al., 2020)* | Figure 5, Figure 6B, Figure 7C-D, Figure S5 |
| 7 | *ProNTP303:YC3.6-ProNTP303:PrpS₁ (Wang et al., 2020)* | Figure 6A and 6C, Figure S6, |

**Table S2. Oligonucleotides, targeting peptide sequences and Golden Gate modules used for transgene construction.** A. List of oligonucleotides used in this study. F, forward primers. R, reverse primers. Bold letters indicate target-binding sequences. B. Targeting peptide sequences fused to roGFP2-Orp1 for subcellular localisation; C. Golden Gate Assembly modules used in this study.

**A**

| Primer name | Sequence (5'-3') |
|---|---|
| *F-A-NTP303* | *TTTTGGTCTCAACCT**GATACACTCGCAACGTGTG**TATCCTAA* |
| *R-B-NTP303* | *TTTTGGTCTCATGTT**GACGTTGTTTTTTTTATT**CTTTAATCCCC* |
| *F-D-NTP303* | *TTTTGGTCTCATCAG**GATACACTCGCAACGTGTG**TATCCTAA* |
| *R-E-NTP303* | *TTTTGGTCTCAGCAG**GACGTTGTTTTTTTTATT**CTTTAATCCCC* |
| *NTP303_F* | *TTGAAGACAA**GGAGATACACTCGCAACGTGTG*** |
| *NTP303_TACT_R* | *TTGAAGACAA**AGTAGACGTTGTTTTTTTTATT*** |
| *NTP303_AATG_R* | *TTGAAGACAA**CATTGACGTTGTTTTTTTTATT*** |
| *Chloro_F* | *TTGAAGACAA**ATACTATGGCTTCTTCTATGCTTTCTTCTGC*** |
| *Chloro_R* | *TTGAAGACAA**CATTGCTCGAACTCTTCCTCCGTTACTAG*** |
| *NLS_F* | *TTGAAGACAA**TACTATGCTGCAGCCTAAGAAGAAGAGAAAGGTTGGAGGAATGTTGTCTTCAA*** |
| *NLS_R* | *TTGAAGACAA**CATTCCTCCAACCTTTCTCTTCTTCTTAGGCTGCAGCATAGTATTGTCTTCAA*** |
| *MTS_F* | *TTGAAGACAAA**TACTATGGCGAGTCGGCGAATCTT*** |
| *MTS_R* | *TTGAAGACAA**CATTTTTGCCACCGTAATCATAGG*** |

**B**

| Targeting compartments | Targeting sequences | References |
|---|---|---|
| Cytoplasm | Nuclear Export Signal (NES) of the protein kinase inhibitor (PKI) from rabbit | Wen et al., 1995; Arnaud et al., 2023 |
| Mitochondria | Mitochondrial targeting signal of the ATPase β subunit from *Arabidopsis thaliana* (98% sequence identity with *Nicotiana plumbaginifolia*) | Sjöling & Glaser, 1998, Carrie et al., 2015 |
| Chloroplasts | Chloroplast transit peptide from the Rubisco small subunit RbcS (synthetic consensus of dicot sequences) | Marillonnet et al., 2004; Engler et al., 2014; Arnaud et al., 2023 |
| Peroxisomes | SKL sequence for targeting to peroxisome | Sparkes et al., 2003; Arnaud et al., 2023 |
| Nuclei | Nuclear localisation signal derived from Simian Virus 40 (NLS SV40) | Kalderon et al., 1984; Arnaud et al., 2023 |

**C**

| Modules used | GoldenGate assembly level | *Modules/constructs produced* | *Reference* |
|---|---|---|---|
| *pICH41233, PCR product ProNTP303* | *0* | *pICH41233-ProNTP303* | *Engler et al., 2014; de Graaf et al., 2012* |
| *pICH41295, PCR product ProNTP303* | *0* | *pICH41295-ProNTP303* | *Engler et al., 2014; de Graaf et al., 2012* |
| *pICH41246, PCR product MTS_F+R* | *0* | *pICH41246-MTS* | *Engler et al., 2014* |
| *pICH41246, PCR product Chloro_F+R* | *0* | *pICH41246-Chl* | *Engler et al., 2014* |
| *pICH47732, pICH41233-ProNTP303, pICH41308-roGFP2-Orp1, pICH41421* | *1* | *ProNTP303:roGFP2-Orp1-NOS* | *Arnaud et al., 2023* |
| *pICH47732, pICH41233-ProNTP303, pICH41246-MTS, pICH41308-roGFP2-Orp1, pICH41421* | *1* | *ProNTP303:MTS-roGFP2-Orp1-NOS* | *Arnaud et al., 2023* |
| *pICH47732, pICH41233-ProNTP303, pICH41246-Chl, pICH41308-roGFP2-Orp1, pICH41421* | *1* | *ProNTP303:Chl-roGFP2-Orp1-NOS* | *Arnaud et al., 2023* |
| *pICH47732, pICH41295-ProNTP303, pICH41308-roGFP2-Orp1-SKL, pICH41421* | *1* | *ProNTP303:roGFP2-Orp1-SKL-NOS* | *Arnaud et al., 2023* |
| *pICH47732, pICH41295-ProNTP303, pAGT707, Oligo NLS_F+R, pICH41308-roGFP2-Orp1-SKL, pICH41421* | *1* | *ProNTP303:NLS-roGFP2-Orp1-NOS* | *Arnaud et al., 2023* |
| *pICH47811, pICH87644, pICH41308-Hygro, pICH44300* | *1* | *Act2:Hygro-TerAct2* | *Arnaud et al., 2023* |
| *pAGM4723, pICH41744, ProNTP303:roGFP2-Orp1-Nos, Act2:Hygro-TerAct2* | *2* | *ProNTP303:roGFP2-Orp1-NOS-Act2:Hygro-TerAct2* | *Arnaud et al., 2023* |
| *pAGM4723, pICH41744, ProNTP303:MTS-roGFP2-Orp1-Nos, Act2:Hygro-TerAct2* | *2* | *ProNTP303:MTS-roGFP2-Orp1-NOS-Act2:Hygro-TerAct2* | *Arnaud et al., 2023* |
| *pAGM4723, pICH41744, ProNTP303:Chl-roGFP2-Orp1-Nos, Act2:Hygro-TerAct2* | *2* | *ProNTP303:Chl-roGFP2-Orp1-NOS-Act2:Hygro-TerAct2* | *Arnaud et al., 2023* |
| *pAGM4723, pICH41744, ProNTP303:roGFP2-Orp1-SKL-Nos, Act2:Hygro-TerAct2* | *2* | *ProNTP303:roGFP2-Orp1-SKL-NOS-Act2:Hygro-TerAct2* | *Arnaud et al., 2023* |
| *pAGM4723, pICH41744, NLS-ProNTP303:roGFP2-Orp1-Nos, Act2:Hygro-TerAct2* | *2* | *ProNTP303:NLS-roGFP2-Orp1-NOS-Act2:Hygro-TerAct2* | *Arnaud et al., 2023* |

**Table S3. Multiple reaction monitoring (MRM) transitions for LC-triple quadrupole MS/MS analysis of TCA cycle acids and glutathione.**

| Compound Name | Quantifier or qualifier ion | Precursor Ion | MS1 Res | Product Ion | MS2 Res | Dwell | Fragmentor (V) | Collision Energy (V) | Cell Accelerator Voltage | Polarity |
|---|---|---|---|---|---|---|---|---|---|---|
| Succinate | Quantifier | 329 | Unit | 91 | Unit | 200 | 135 | 20 | 4 | Positive |
| Succinate | Qualifier | 329 | Unit | 206 | Unit | 200 | 135 | 20 | 4 | Positive |
| Malate | Quantifier | 345 | Unit | 91 | Unit | 200 | 135 | 20 | 4 | Positive |
| Malate | Qualifier | 345 | Unit | 222 | Unit | 200 | 135 | 20 | 4 | Positive |
| 2-Oxoglutarate | Quantifier | 462 | Unit | 91 | Unit | 200 | 135 | 20 | 4 | Positive |
| 2-Oxoglutarate | Qualifier | 462 | Unit | 339 | Unit | 200 | 135 | 20 | 4 | Positive |
| Aconitate | Quantifier | 490 | Unit | 91 | Unit | 200 | 135 | 20 | 4 | Positive |
| Aconitate | Qualifier | 490 | Unit | 325 | Unit | 200 | 135 | 20 | 4 | Positive |
| Citrate_Isocitrate | Quantifier | 508 | Unit | 91 | Unit | 200 | 135 | 20 | 4 | Positive |
| Citrate_Isocitrate | Qualifier | 508 | Unit | 385 | Unit | 200 | 135 | 20 | 4 | Positive |
| Glutathione (GSH) | Quantifier | 413 | Unit | 91 | Unit | 200 | 135 | 20 | 4 | Positive |

### References

Arnaud D, Deeks MJ, Smirnoff, N. Organelle-targeted biosensors reveal distinct oxidative events during pattern-triggered immune responses. *Plant Physiology*. 2023:191(4):2551–2569. https://doi.org/10.1093/plphys/kiac603

Carrie C, Venne AS, Zahedi, RP, Soll J. Identification of cleavage sites and substrate proteins for two mitochondrial intermediate peptidases in *Arabidopsis thaliana*. *J Exp Bot*. 2015:66(9):2691-2708. https://doi.org/10.1093/jxb/erv064

de Graaf BH, Vatovec S, Juarez-Diaz JA, Chai L, Kooblall K, Wilkins KA., Zou H, Forbes T, Franklin FC, Franklin-Tong VE. The *Papaver* self-incompatibility pollen *S*-determinant, PrpS, functions in *Arabidopsis thaliana*. *Curr Biol*. 2012:22(2):154-159. https://doi.org/10.1016/j.cub.2011.12.006

Engler C, Youles M, Gruetzner R, Ehnert TM, Werner S, Jones JD, Patron NJ, Marillonnet S. A golden gate modular cloning toolbox for plants. *ACS Synth Biol.* 2014:3(11):839-843. https://doi.org/10.1021/sb4001504

Kalderon D, Richardson WD, Markham AF, Smith AE. Sequence requirements for nuclear location of simian virus 40 large-T antigen. *Nature.* 1984:311(5981):33-38. https://doi.org/10.1038/311033a0

Marillonnet S, Giritch A, Gils M, Kandzia R, Klimyuk V, Gleba Y. In planta engineering of viral RNA replicons: efficient assembly by recombination of DNA modules delivered by *Agrobacterium*. *Proc Natl Acad Sci U S A*. 2004:101(18):6852-6857. https://doi.org/10.1073/pnas.0400149101

Sjöling S, Glaser E. Mitochondrial targeting peptides in plants. *Trends in Plant Science.* 1998:3(4):136-140. https://doi.org/10.1016/S1360-1385(98)01212-6

Sparkes IA, Hawes C, Baker A. AtPEX2 and AtPEX10 are targeted to peroxisomes independently of known endoplasmic reticulum trafficking routes. *Plant Physiol. 2005:139*(2):690-700. https://doi.org/10.1104/pp.105.065094

Wen W, Meinkoth JL, Tsien RY, Taylor SS. Identification of a signal for rapid export of proteins from the nucleus. *Cell.* 1995:82(3):463-473. https://doi.org/10.1016/0092-8674(95)90435-2
